# Statistical power and false positive rates for interdependent outcomes are strongly influenced by test type: Implications for behavioral neuroscience

**DOI:** 10.1101/2023.05.01.538922

**Authors:** Michelle Frankot, Peyton M. Mueller, Michael E. Young, Cole Vonder Haar

## Abstract

Statistical errors in preclinical science are a barrier to reproducibility and translation. For instance, linear models (e.g., ANOVA, linear regression) may be misapplied to data that violate assumptions. In behavioral neuroscience and psychopharmacology, linear models are frequently applied to interdependent or compositional data, which includes behavioral assessments where animals concurrently choose between chambers, objects, outcomes, or types of behavior (e.g., forced swim, novel object, place/social preference). The current study simulated behavioral data for a task with four interdependent choices (i.e., increased choice of a given outcome decreases others) using Monte Carlo methods. 16,000 datasets were simulated (1,000 each of 4 effect sizes by 4 sample sizes) and statistical approaches evaluated for accuracy. Linear regression and linear mixed effects regression (LMER) with a single random intercept resulted in high false positives (>60%). Elevated false positives were attenuated in an LMER with random effects for all choice-levels and a binomial logistic mixed effects regression. However, these models were underpowered to reliably detect effects at common preclinical sample sizes. A Bayesian method using prior knowledge for control subjects increased power by up to 30%. These results were confirmed in a second simulation (8,000 datasets). These data suggest that statistical analyses may often be misapplied in preclinical paradigms, with common linear methods increasing false positives, but potential alternatives lacking power. Ultimately, using informed priors may balance statistical requirements with ethical imperatives to minimize the number of animals used. These findings highlight the importance of considering statistical assumptions and limitations when designing research studies.

## Introduction

Translation of preclinical research is a major challenge with estimates that up to 90% of therapeutics in behavioral neuroscience fail to translate to humans [1]. While some causes cannot be easily remedied, statistical analysis and its misapplication can be readily addressed. Reviews in the fields of pharmacology and neurotrauma found that up to 84% and 70% of papers, respectively, contained inappropriate statistics or inference [2, 3]. The use of parametric tests when core assumptions are violated can produce inaccuracies, increase false positives (Type I error), and reduce power (Type II error). Violations may often go unnoticed; when 30 PhD researchers were given hypothetical datasets to analyze, fewer than 25% checked assumptions [4]. For linear models (e.g., t-test, ANOVA, linear regression), the residuals (i.e., the difference between observed and predicted values) must meet several assumptions, including that they are independent of one another [5].

Many behavioral assays in neuroscience can produce data that violate the independence assumption, particularly when considering how those data are commonly analyzed. Numerous tasks allow animals to choose between different chambers, objects, outcomes, or types of behavior to engage in. While two-choice tasks (e.g., novel object recognition, elevated plus maze, two-lever operant choice tasks) present a simple solution of analyzing a single outcome, other behaviors have multiple outcomes of interest. For example, in the Forced Swim Task, a measure of behavioral despair, rodents float after initial swimming and escape behaviors [6]. When considered together for analysis, these behaviors are compositional in nature and interdependent because an increase in one inherently decreases the others (i.e., they sum to 100%). Interestingly, at least some antidepressants mediate switches to specific behaviors (e.g., from floating to swimming or floating to escaping) [7]. While the switch *away* from floating may be sufficient to draw conclusions regarding depressive-like behavior, for other tasks this is likely not the case. Three-chamber tasks (e.g., social preference, conditioned place preference) are quite popular because of the ability to dissociate preference, aversion, and indifference [8], but suffer the same issues.

In analyzing such data, it is relatively common practice to aggregate (i.e., sum, average) across each option and analyze using linear models for several reasons (e.g., ease of interpretation, familiarity). Analysis of interdependent outcomes in such paradigms clearly violate assumptions for linear models and an analysis method should account for this [9]. It is well-known that analysis of non-aggregated data using generalized linear models (GLMs) or multivariate analyses will largely address account for this compositional nature [10]; however, barriers such as computational demand and a steeper learning curve make these less accessible. Despite the common application of linear models to interdependent, compositional aggregated data, we do not currently know the extent to which this will impact conclusions drawn from significance testing. Linear models are sometimes accurate despite violations [11], although there is debate on this topic [12-14]. Thus, it is important to empirically determine the accuracy of various approaches. To do so, statistical methods can be evaluated via Monte Carlo simulation, which involves repeated sampling of random observations from population parameters (e.g., effect size) set by the researcher. For example, such simulations showed that mixed-effect modeling and censored regression were more accurate than ANOVA for Morris water maze data [15, 16].

To determine which models should be assessed for accuracy in a simulation study, we considered both common methods and those which may best account for interdependent data. The linear models (often on aggregated data) may be the most common analyses in neuroscience [17] likely because they are covered in undergraduate and graduate statistics courses as a minimum requirement. However, analysis of the trial level using a binomial or multinomial logistic regression can account for the interdependence of data and may increase power. Many researchers have also come to consider mixed effects (also called multi-level, hierarchical, random effects) models because they account for nested data (e.g., multiple observations within a subject), handle missing data, and provide additional information about within-group variance [18]. Mixed models can be applied to both linear and nonlinear analyses. Finally, Bayesian versions of these models, which incorporate prior belief (e.g., informed by historical data) may further augment power as has been recently demonstrated for t-tests [19].

In the current study, we demonstrate via simulation that small changes to statistical methodology can have a profound impact on accuracy when analyzing interdependent data. To evaluate the accuracy of these statistical methods for interdependent data, we applied Monte Carlo techniques to simulate concurrent 4-choice data. Our simulation modeled real behavioral data from the Rodent Gambling Task (RGT) [20] with parameters derived from a publicly-available dataset [21]. 1,000 datasets were simulated for each of four sample sizes and four effect sizes (N = 16,000 datasets). We then compared the accuracy of common linear models which neglect interdependencies. These results were then contrasted against other potential approaches to handle interdependency, including an aggregate score, random effects for each option, and logistic regressions. In our last analysis, we set priors for control subjects in a Bayesian analysis to increase power. Finally, we validate these findings against a novel dataset collected in a different laboratory.

## Materials and Methods

### Overview

To model interdependency, RGT data were simulated and analyzed using R statistical software (https://www.r-project.org/). Simulations reflected actual rodent behavior collected from multiple studies and included control subjects (sham surgery: “Control”), subjects with a brain injury (traumatic brain injury: “TBI”), a second set of control subjects (normal RGT: “Control”), and subjects exposed to complex audiovisual cues during reinforcement (cued RGT: “Cue”).

### Rodent Gambling Task (RGT)

The task was conducted in a standard operant chamber with a 5-hole array, with four options presented in each trial (Figure 1A) [20]. In brief, rats were initially trained to respond to a randomly illuminated hole until they could respond during a 10 s illumination. A requirement to inhibit responses until lights were illuminated (typically 5 s from trial initiation) was used as a measure of motor impulsivity and punished with a 5 s timeout. Omitted responses were also punished with a 5 s timeout. Once trained, rats were given 7 forced-choice sessions of the RGT, which ensured options were sampled equally. Then, rats were allowed to freely choose for daily 30-min sessions. In the full RGT task, array lights were illuminated at the beginning of each trial, and a response turned off the lights and resulted in reinforcement (sucrose pellets) or punishment (timeout from reinforcement). The 2-pellet option is “optimal” because it has the highest average reinforcement rate (80% chance of 2 pellets, 20% chance of 10 s time out). The 1-pellet option is considered “suboptimal” (90% chance of 1 pellet, 10% chance of 5 s time out), and the 3- and 4-pellet options are “risky” because they often result in lengthy timeouts and lower rates of reinforcement (3-pellet: 50% chance of 3 pellets, 50% chance of 30 s time out; 4-pellet: 40% chance of 4 pellets, 60% chance of 40 s time out). The location of the holes was counterbalanced across animals to account for potential side bias.

**Figure 1.**
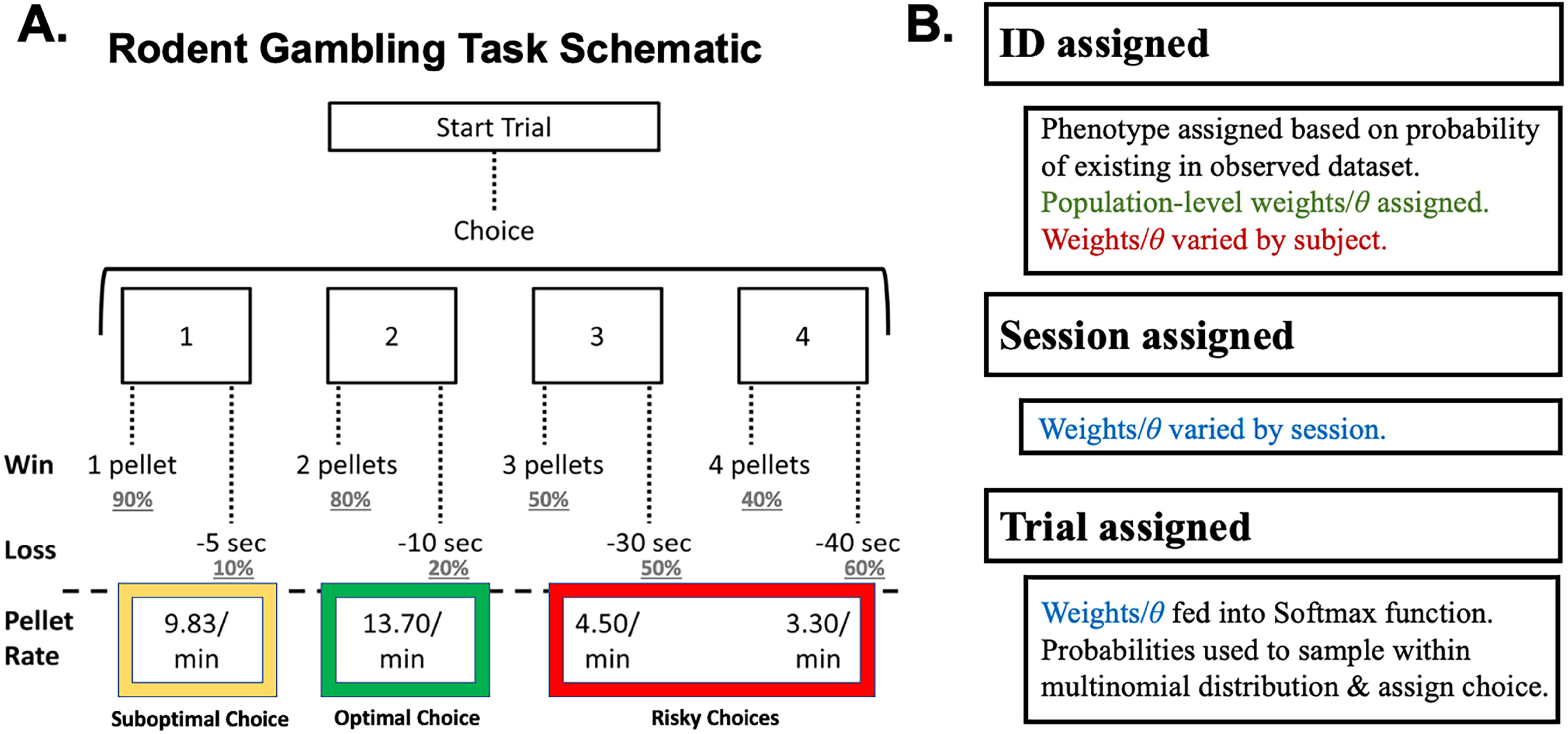
Task and simulation design. A) Schematic of the Rodent Gambling Task [51]. After initiating a trial, rats chose from any of the four holes. Each hole was associated with a different probability and magnitude of reinforcement and punishment. As a result of varying reinforcement rates, the 1-pellet option (P1) was suboptimal, the 2-pellet option (P2) was optimal, and the 3- and 4-pellet options (P3, P4) were risky. B) Logical flow of the simulation, where a subject is assigned a phenotype and softmax parameters. Those parameters were varied slightly between subjects and across sessions and then used to generate a choice (optimal, suboptimal, or risky) on each simulated RGT trial.

This task generates data at an individual trial level: the choice is recorded for each trial. Omitted or premature responses and various latencies are also recorded, but are irrelevant for the current analyses. Each individual trial is independent from the former outcomes, but animals must choose only one option on a given trial. Changes in preference of one option over others is then quantified based on individual subjects and experimental manipulations. Data is output in a trial by choice matrix and frequently aggregated at the session level to a compositional format: percent choice of each option.

### Datasets

A publicly-available dataset of behavior on this task [21] collected by our laboratory, which we refer to as the “control set” was used to inform simulation parameters. For the current study, only stable post-injury data (i.e., collected outside the initial task-learning and acute injury phases) were considered, and data involving manipulations (e.g., treatments) other than brain injury were excluded. This resulted in 109 subjects (*n* = 51 for TBI; *n* = 58 for Control). The original studies from which these data were acquired were approved by the West Virginia University Institutional Animal Care and Use Committee.

A second dataset assessing this task, which we refer to as the “validation set” was obtained from Dr. Catharine Winstanley and is fully described elsewhere [22]. In brief, this dataset consisted of 14 studies of rats learning the RGT. We selected sessions above 15 (resulting in 5-10 sessions per study) to more closely approximate stable behavior used in the control set. 328 total subjects were in either normal condition (i.e., identical to the control set) or “cued” condition, where complex audiovisual cues were paired with pellet delivery (n = 168 for Control, n = 160 for Cued). The original studies from which these data were acquired were approved by the University of British Columbia Animal Care Committee. This was selected as a validation set because the manipulation generates a distinct effect from that of TBI (i.e., biases rats toward the 3- and 4-pellet options).

### Monte Carlo Simulation

Code for generating these data and the generated data is provided in the supplemental material (S2, Simulations.zip). The current simulations approximated stable, learned behavior to focus analyses on the effects of interdependency and most closely replicate the data they were generated from. Data were generated at the individual trial/choice level for individual simulated subjects and for 10 sessions of data to closely mimic real-world conditions on this behavior. No within- or between-session learning effect was modeled. Choice data were generated at the trial level by fitting the control set to a 2-parameter softmax function, which takes in a list of values and returns probabilities: 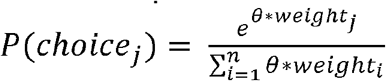, where *j* (1 to 4) is the index of the choice option, *weight*_*j*_ is equal to average value of the *j*^th^ choice, and θ is equal to the degree of exploration among options.

For the control set, we previously identified five unique behavioral phenotypes on this task using *k*-means clustering [23]. Thus, we fit the softmax function to each phenotype separately, to generate *weight* and θ values as well as their standard deviations that were used as population parameters. Each simulated subject was assigned a phenotype (θ and *weight* values) based on the probability of that phenotype existing in the control set. Then, softmax parameters were sampled from the population for that phenotype and varied between subjects according to the standard deviation and across 10 sessions. Parameters were passed through the softmax function, producing probabilities that were used to generate discrete choices at the trial level for each subject. Prior to generating large datasets, the simulated data were evaluated against real data using aggregate descriptions (i.e., mean, standard deviation) and visual inspection to confirm that simulated individual subjects were representative of common and rare actual subjects (see Figure S1 for an example).

The simulation was conducted in a nested for-loop structure to automate dataset generation (Figure 1B). 1,000 datasets were simulated at four sample sizes (*n*/group = 6, 10, 14, 20) and four effect sizes representing the effect of TBI on optimal choice (Cohen’s *f* = 0.0, 0.3, 0.4, 0.5). The *f* = 0.4 effect size is representative of the TBI effect in the control set.

For the validation set, a similar approach was taken. The effect of cue exposure was substantially different than TBI: cues selectively increased choice of risky options [24]. We identified five behavioral phenotypes with k-means clustering, some of which overlapped with the control set and some which were distinct, and used those for simulation. The magnitude of the cue exposure effect size was similar at *f* = 0.37, so 0.4 was used for the most possible direct comparison with the control set. 1000 datasets at *f* = 0.4 and *f* = 0 were simulated for sample sizes of 6, 10, 14, and 20 per group.

### Data Analysis

The strategy for model selection was based upon what is commonly used in the field of behavioral neuroscience and models that should best account for these types of data. As such, a linear, fixed effect only model was provided as an initial point of comparison due to its ubiquity. Subsequent models incorporated mixed effects to account for inter-subject variance and allow for more precise parameter estimation as has been previously demonstrated [15].

Datasets were analyzed in an automated fashion, where a regression was conducted for the first dataset, results were written to a file, and a regression was conducted on the next set until completion (see Figure 2). The accuracy of common linear models was tested in Experiment 1, methods accounting for interdependency in Experiment 2, and the effects of Bayesian priors in Experiment 3. Any reported errors or warnings were recorded to a table for review. A small number of models for M4 (∼20%) had convergence warnings at the default *lme4* settings, likely due to the nature of the random effects accounting for substantial variance and limiting iterations. There were also some minor, not uncommon warnings in the Bayesian models identifying 1-2 divergent transitions (at low N sample size), and treedepth warnings (at higher sample size). A very small number warned about effective sample size (ESS; on the 0 effect size simulations). Assumptions were checked visually on a subset of models (check_model() function from *performance* package). There were minor deviations to homoscedasticity which were common across multiple model types, suggesting it is not a major driver of any reported effects.

**Figure 2.**
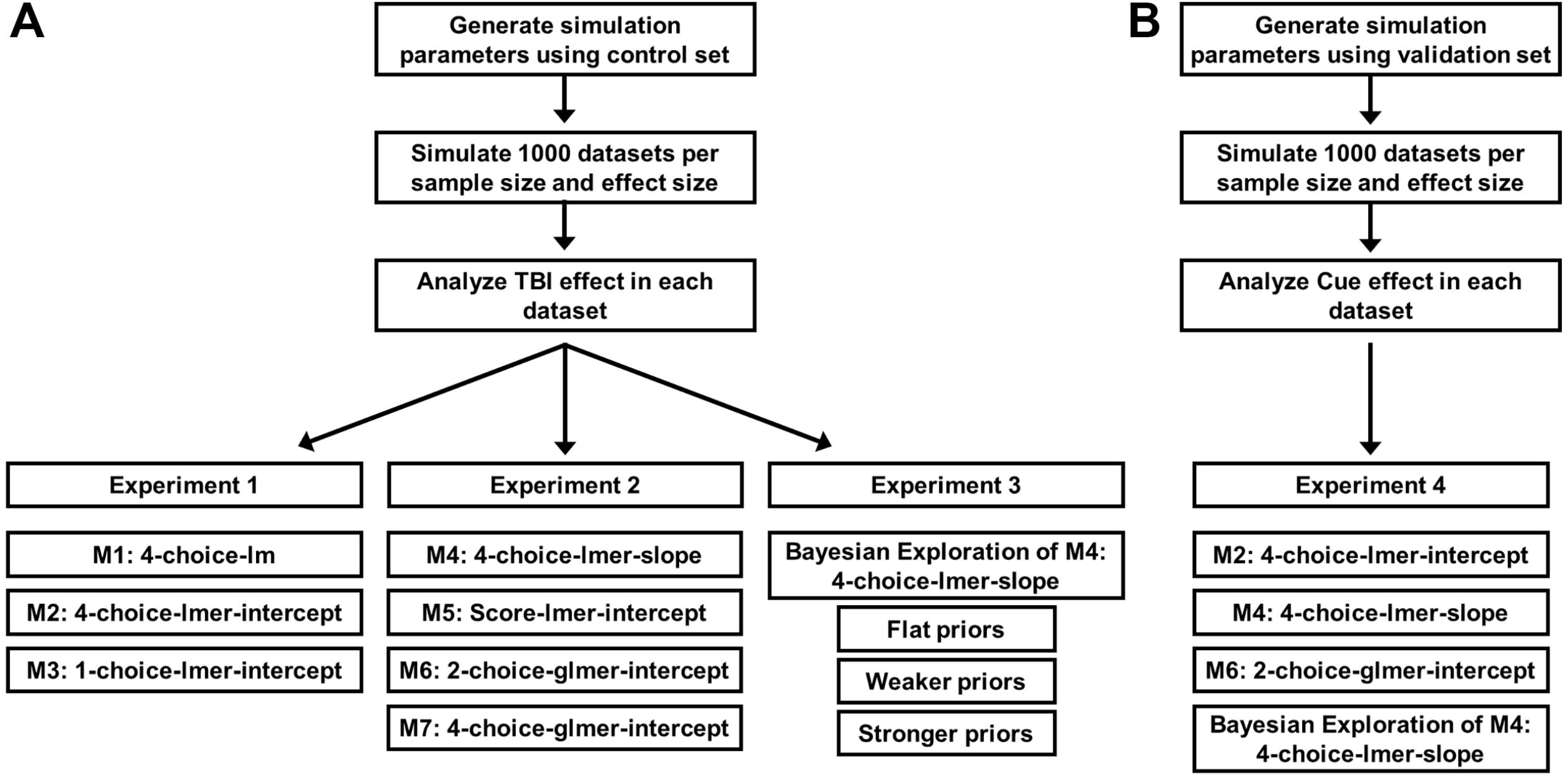
Experimental design. A) Simulation parameters were generated and used to create 16,000 datasets (4 sample sizes and 4 effect sizes) with Control (sham surgery) and TBI (traumatic brain injury) rats. A series of three experiments were conducted to compare the accuracy of different statistical approaches. B) A fourth experiment generated new data (8000 datasets; 4 sample sizes and 2 effect sizes) from a second dataset assessing Control (regular task) and Cued (audiovisual cue task) rats to evaluate the key findings from the first three experiments.

Mixed models were performed using the *lme4* package [25], with pseudocode for each model provided below. Significance was determined via model comparison (i.e., comparing models with and without TBI via likelihood ratio test). When TBI improved model fit at *p* < 0.05, the injury effect was considered significant. Four outcomes were possible: true positive (TP; significant effect when *f* > 0), false positive (FP; significant effect when *f* = 0), false negative (no significant effect when *f* > 0), and true negative (no significant effect when *f* = 0). *Z*-scored TP and FP rates were used to calculate the signal detection theory metrics of accuracy/discriminability (*d*’; *z*(TP) - *z*(FP)) and bias (*c*; -½*[*z*(TP) + *z*(FP)]) [26]. Variance-covariance matrices in random effects were generated by the model specification: for most models, an unstructured intercept-only matrix was used. Only in M4 was a variance-covariance matrix constructed between slope and intercept terms.

### Experiment 1

In Experiment 1, we compared the accuracy of three linear models which do not account for interdependencies but are commonly used to analyze such data. An arcsine-square root transformation was applied to improve the normality of the error term distribution, as is common convention. First, choice of all four outcomes was analyzed concurrently in a fixed effects regression, analogous to common linear regression or ANOVA (“M1: 4-choice-lm”; Percent choice as a function of choice option, injury, and their interaction). Second, all four choices were again analyzed concurrently, but in a mixed effects model and the intercept (dummy coded as “optimal” choice) was varied by subject as the random effect (“M2: 4-choice-lmer-intercept”; Percent choice as a function of choice option, injury, and their interaction with initial preference for choice option 2 as a random effect at the subject level). Lastly, data was reduced to one choice only (“optimal” option) and analyzed using a linear model with the intercept as the random effect (“M3: 1-choice-lmer-intercept”; Percent choice of option 2 as a function of injury with initial preference for option 2 as a random effect at the subject level) which would ultimately need to be repeated four times (once for each option) with each analysis affected by interdependency. Below is R pseudocode for reference:

- M1 4-choice-lm: lm(PctChoice∼Option*Injury)
- M2 4-choice-lmer-intercept: lmer(PctChoice∼Option*Injury + (0+dummy(Option, “2”)|Subject)) #note that the random effects term must be dummy coded manually in R’s *lme4*
- M3 1-choice-lmer-intercept: lmer(PctChoice_Optimal∼Injury + (1|Subject))

### Experiment 2

In Experiment 2, we accounted for interdependency by modifying M2 above to allow all choices to vary by subject in the random effect in a 4-choice mixed effect model (“M4: 4-choice-lmer-slope”; Percent choice as a function of option, injury, and their interaction with preference for each option as a random effect at the subject level). Next, we determined the viability of a score variable, a common workaround to avoid analyzing concurrent outcomes. Risky choices were subtracted from optimal+suboptimal choices to form a score and assessed in a mixed effect model with subject-level intercepts in the random effect (“M5: Score-lmer-intercept”; Score as a function of choice option, injury, and their interaction with the random effect of initial score at the subject level). Next, we considered whether reducing data similar to M3 above, but in a fashion which accounts for dependence, would more accurately model choice. We analyzed optimal versus other outcomes in a 2-choice binomial mixed effects logistic regression (generalized linear mixed effects regression with logit link function; “M6: 2-choice-glmer-intercept”; Proportion choice as a function of injury with initial choice preference as a random effect at the subject level). Again, a drawback of this method is a need to repeat for multiple comparisons. To address this limitation, we also analyzed all four choices concurrently in a Bayesian 4-choice multinomial logistic regression (“M7: 4-choice-glmer-intercept”; flat priors; Proportion choice as a function of choice option, injury, and their interaction with initial choice preference as a random effect at the subject level). This analysis required the *brms* package [27]. Because Bayesian analyses do not provide a significance test, an effect was considered “significant” if the credible interval for the TBI effect on optimal versus suboptimal choice did not contain zero. This approach was used instead of model comparison due to the computational intensity of running multiple multinomial models. Below is R pseudocode for reference:

- M4 4-choice-lmer-slope: lmer(PctChoice∼Option*Injury + (1+Option|Subject))
- M5 Score-lmer-intercept: lmer((Optimal – Risky)∼Injury + (1|Subject))
- M6 2-choice-glmer-intercept: glmer(Optimal/Total∼Injury + (1|Subject), weights=Total, family = “binomial”)
- M7 4-choice-glmer-intercept: brm(Optimal/Total | Option∼Injury + (1|Subject), weights=Total, family= “categorical”)

### Experiment 3

A third experiment was conducted to increase power using Bayesian inference. We used M4, the 4-choice random slope model for several reasons: (1) accuracy was similar to other best-performing models, (2) the R implementation might be familiar to researchers, (3) the results are more readily interpretable than GLMs, and (4) it concurrently analyzes all four outcomes, removing the need for multiple comparisons and corrections.

In Bayesian analyses, prior beliefs about the parameter distribution are integrated with new experimental data to produce a posterior distribution, which provides the most likely parameter values (for a beginner-friendly review, see [28]). Prior beliefs may be specified from general knowledge about the data-generating process, derived from previous experimental results, or assumed to be uninformed (“flat” priors). Priors for Control rats only (means and standard deviations derived from a 4-choice random slope model fit on the control set from which simulated data were derived) were used to increase power as per Bonapersona et al. [19].

We compared three models: one with flat priors, a second with weaker priors (exact means, inflated error; i.e., a wider prior distribution), and a third with strong priors calculated from the control set (exact parameters of the control set; i.e., a narrow prior distribution). Although Bayesian implementations recommend against arbitrary cut-offs (e.g., *p*-values), for ease of comparison across models, we used the *brms::hypothesis* function for significance testing. If the 95% credible interval for the TBI effect on optimal choice did *not* contain zero, the effect was considered significant. These models were performed on a random sample of 100 datasets at an effect size of *f* = 0.0 and 0.4 (our observed TBI effect from real data) across samples sizes of *n* = 6, 10, and 14 per group. The remaining datasets were not analyzed due to computation time.

To determine whether changing to Bayesian analysis affected previously-published data, we re-analyzed three existing publications using our weaker priors specified above (see supplement S1 for details).

### Experiment 4

To validate the findings of the prior experiments, we replicated the process of generating and analyzing data using the validation set, which included a different experimental manipulation (“cued” versus “control”) and generated a different type of effect. We assessed one problematic model from Experiment 1 (M2), our accurate but underpowered models from Experiment 2 (M4 and M6), and the Bayesian implementation of M4 from Experiment 3 at effect sizes of *f* = 0 or 0.4. The Bayesian analyses used a random subset of 100-200 datasets. Priors were extracted from a model fit of the population data on which the simulation is based and the error term was matched to the original control set for more direct comparisons of outcome.

## Results

### Experiment 1

The results for M1 (4-choice-lm) and M2 (4-choice-lmer-intercept) models followed a similar pattern. Both demonstrated exceedingly high false positive rates above the 5% target (Figure 3A; >70% and >60%, respectively) alongside very low false negative rates (Figure 3B). Bringing these together in a signal detection theory analysis showed that these models are not appropriate for interdependent data as reflected by poor discriminability (Figure 3C) and bias toward positive results (Figure 3D). The M3 (1-choice-lmer-intercept) model had a more reasonable false positive rate ranging from 4-6% across sample sizes (Figure 3A). However, power of this model was limited; even a sample size of 20 did not achieve 80% power at our real-world TBI effect (*f* = 0.4; Figure 3B). Both large effects and samples would be required for sensitivity to effects without bias for M3 (Figure 3C-D). Thus, additional models were explored in Experiment 2.

**Figure 3.**
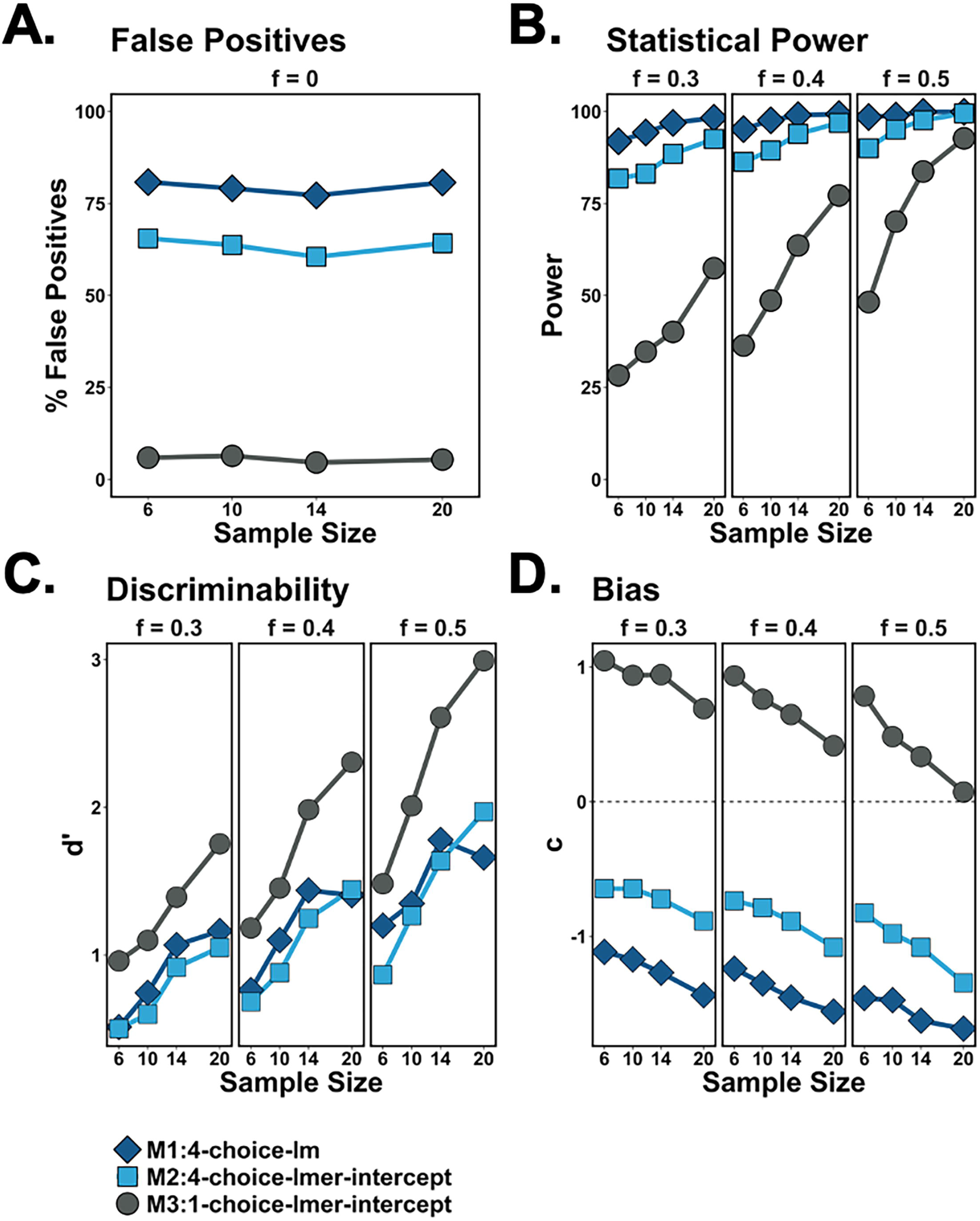
Results of Experiment 1. False positives (Type I error; Panel A), power (1-Type II error; Panel B), discriminability (Panel C), and bias (Panel D) were quantified across three linear models. None of the models accounted for interdependency in the data. M1 (dark blue diamonds) and M2 (light blue squares) performed poorly and were heavily biased toward false positives. M3 (gray circles) better discriminated between outcomes but had low power and would require three additional analyses to understand all effects.

### Experiment 2

All models in Experiment 2 had reasonable false positive rates that varied slightly around 5% (Figure 4A). In terms of power, M5 (Score-lmer-intercept) model performed poorly, never reaching a desirable 80% power. The other models were more reasonably powered, but only the M7 (4-choice-glmer-intercept) model obtained 80% power at N = 20 for the real-world effect size of *f* = 0.4. The M4 (4-choice-lmer-slope) and M6 (2-choice-glmer-intercept) models were able to reach this threshold with a larger effect size (*f* = 0.5), but not until N equaled 20 or 14 per group, respectively (Figure 4B). Signal detection analyses indicated that for M4, M6, and M7 models, discriminability was still lower than desired at a real-world effect and sample sizes (Figure 4C, middle panel) and conservatively biased against detecting an effect (Figure 4D). Based on these findings, we explored whether setting priors for Control subjects in M4 would improve accuracy in Experiment 3.

**Figure 4.**
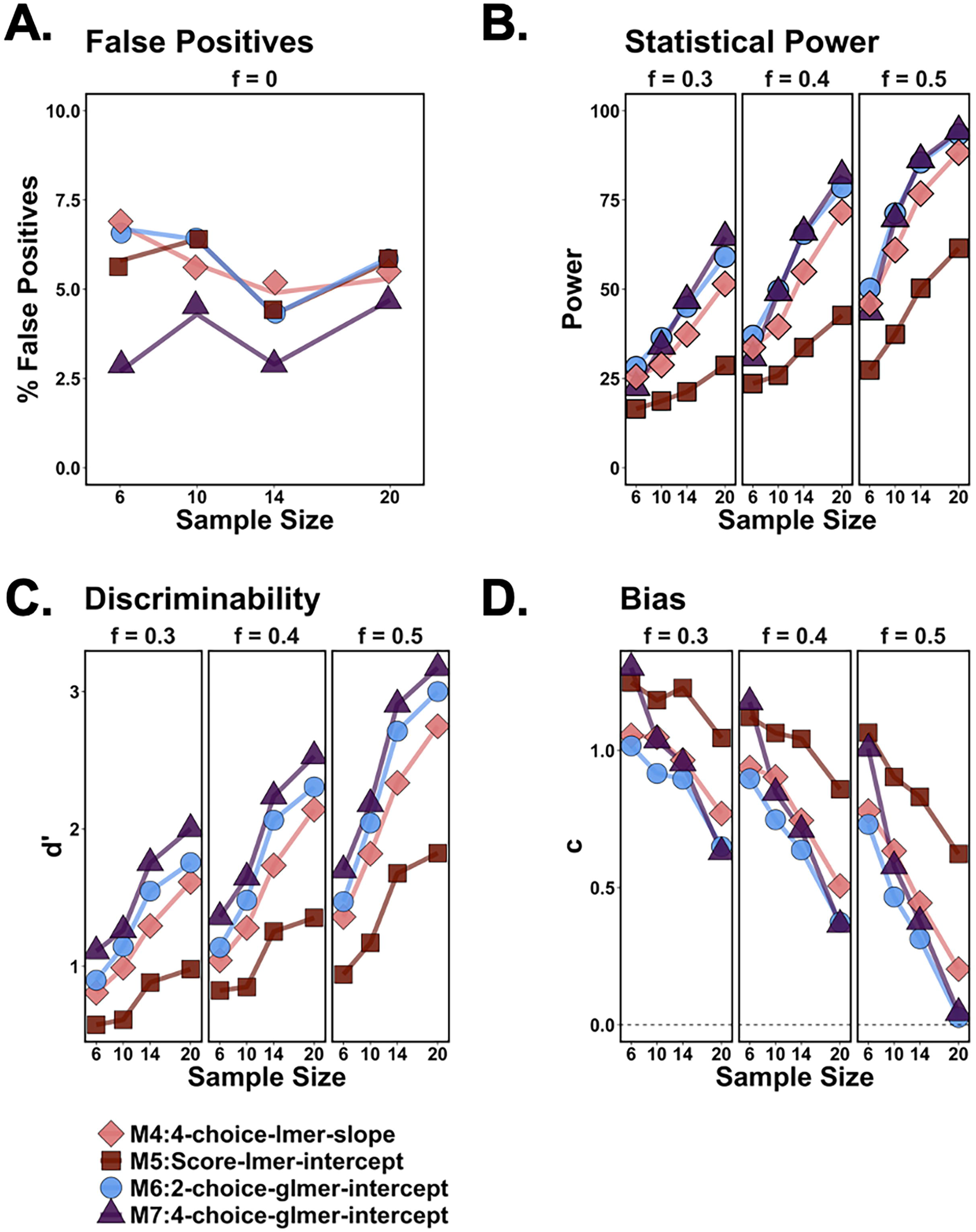
Results of Experiment 2. False positives (Type I error; Panel A), power (1-Type II error; Panel B), discriminability (Panel C), and bias (Panel D) were quantified across four models. Each model accounted for interdependency in some fashion. Lines and points were slightly jittered for visualization. M5 (dark red squares) was underpowered compared to the other models. M4 (pink diamonds), M7 (light blue circles), and M8 (purple triangles) all varied around 5% false positives and were less conservative than the score variable model in detecting real effects. However, these models required large effect and sample sizes to approach or exceed 80% power.

### Experiment 3

Setting priors on Control subjects had minimal effects on false positives, with the potential exception of the larger sample size (Figure 5A). However, priors drastically increased power, graded by strength of the prior (Figure 5B). At a sample size of *n* = 6 per group and Cohen’s *f* = 0.4 (real-world TBI effect), priors increased power from 44% to 69% or 83% for the weak and strong priors, respectively. Results were similar at sample sizes of *n* = 10 and 14 per group. Likewise, discriminability increased with strength of prior and sample size (Figure 5C). Conservative bias (positive direction; toward no effect detected) was also reduced by informed priors, but strong priors at a larger sample size was biased toward positive results (Figure 5D). Ultimately, the weak and strong informed prior models exceeded performance of all other models described in Experiments 1 and 2. Applying weak priors to previously-published data affected the interpretation of TBI effects over time but did not change the overall conclusions of the studies (see Supplement S1 and Table S1 for details). Flat priors were even effective in improving the power of the similarly-specified M4 (n = 6: +10.3%, n = 10: +20.5%, n = 14: +7.1%).

**Figure 5.**
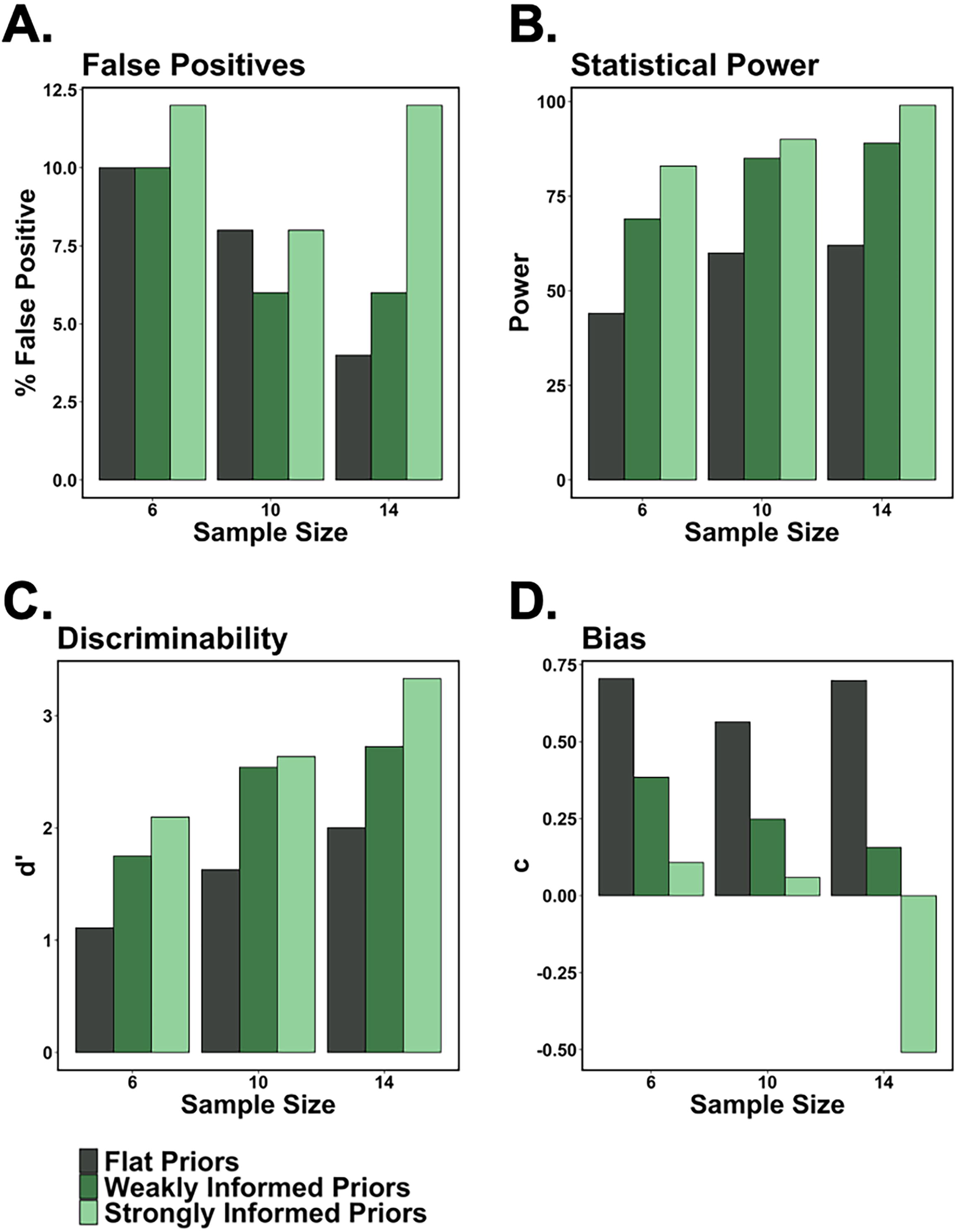
Results of Experiment 3. False positives (Type I error; Panel A), power (1-Type II error; Panel B), discriminability (Panel C), and bias (Panel D) were quantified across three different Bayesian models for 100 datasets per sample/effect combination. The model with flat priors (dark gray bars) was underpowered and the lowest performing in terms of discriminability. Weakly informed (dark green bars) and strongly informed priors (light green bars) improved power, discriminability, and reduced bias. However, strongly informed priors also increased false positives and biased toward detection at the highest sample size.

### Experiment 4

Even when considering a different experimental manipulation of Cued versus Control, the same analysis patterns emerged. M2 still grossly inflated false positive rates (Figure 6A) to slightly under 60% across all sample sizes. M4 and M6 rescued false positive problems, but only approached 70% power at the *n* = 20 per group sample size (Figure 6B), suggesting they will still discriminate poorly and be biased toward non-detection at a real-world effect size of *f* = (Figure 6C-D). The Bayesian implementation of M4 improved power with flat (11%), weak (26-30%), and strong priors (35-42%; Figure 6F), however false positive rates were slightly elevated at closer to 10% than the target 5% (Figure 6E). Despite this, they still discriminated well and reduced non-detection bias (Figure 6G-H).

**Figure 6.**
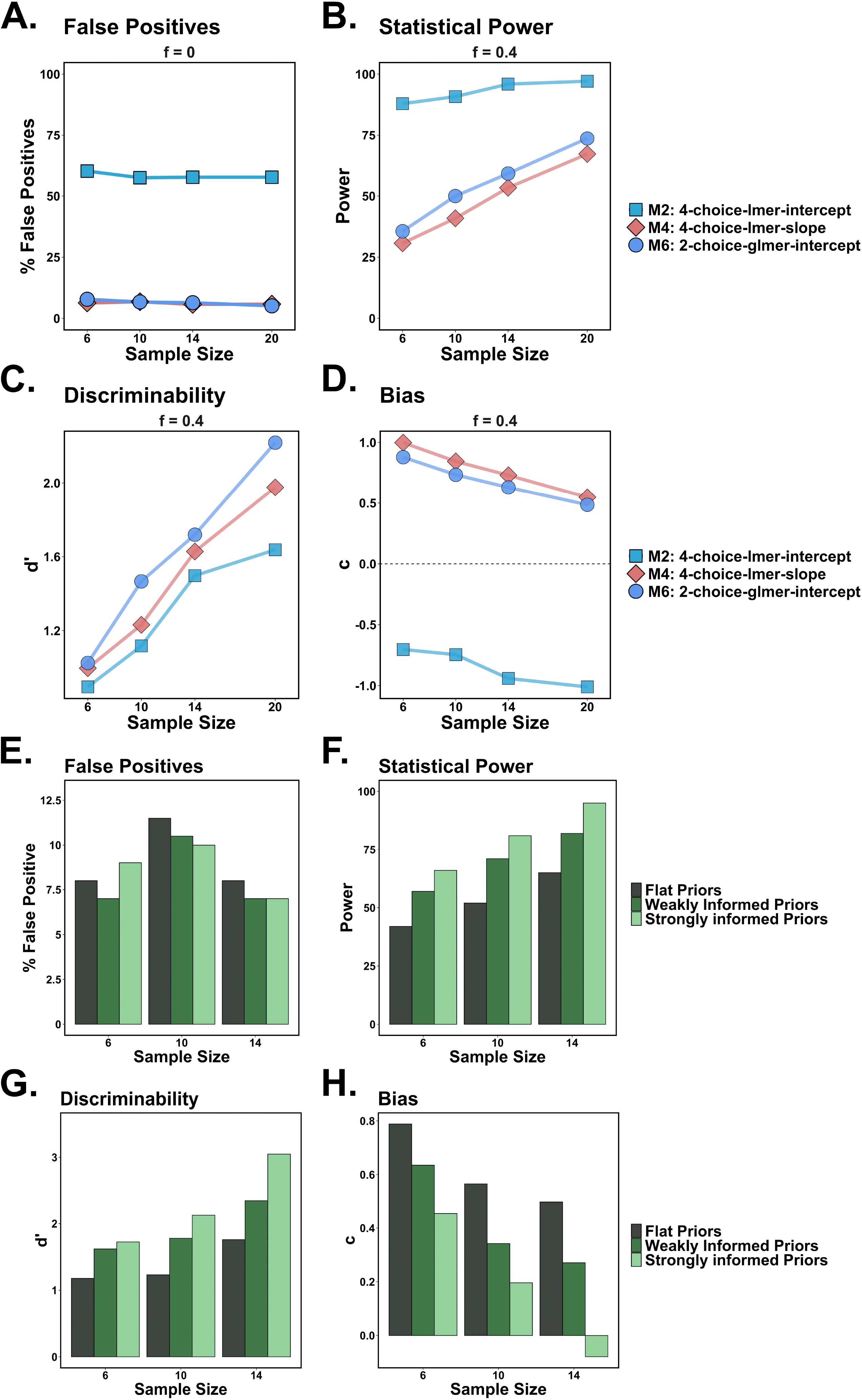
Results from Experiment 4 on the validation data set. False positives (Type I error; Panel A, E), power (1-Type II error; Panel B, F), discriminability (Panel C, G), and bias (Panel D, H) were quantified across models. Panels A-D represent comparisons of M2, M4, and M6 at the *f* = 0.4 effect size. These largely map onto the findings from prior experiments with regard to high false positives in the interdependent linear model (M2), and relatively low power in models that did account for dependency (M4, M6). Panels E-H assessed Bayesian models using different prior belief parameters. Stronger priors increased power and discriminability, similar to findings in Experiment 3, however false positive rates were slightly elevated, although this did not substantially bias results towards toward detection.

## Discussion

Reliability and sensitivity of measures is crucial for strong scientific inquiry. The use of inappropriate statistical models may contribute to problems in replication and the failure to translate from animal models to human populations [29]. In the current study, we simulated interdependent choice data and a biological manipulation (brain injury). Commonly-used fixed-effect and mixed intercept-only linear models were both highly biased toward false positives (Experiment 1; Figure 3). Logistic and linear models that allowed each choice to vary by subject reduced bias toward positives but were underpowered at common preclinical sample sizes (Experiment 2; Figure 4). We substantially increased power by applying priors to control subjects in a Bayesian analysis (Experiment 3; Figure 5) and confirmed all these effects in a second, novel set of simulations (Experiment 4; Figure 6). These results highlight the importance of model selection/specifications when analyzing interdependent data and demonstrate the benefits of Bayesian inference when larger sample sizes are impractical. Additional resources and expanded discussion can be found in supplement S1.

Appropriate model selection is crucial for accurate statistics. Approaches are commonly informed by tradition [30], and reported methods often lack justifications [31]. This may be exacerbated in neuroscience given that statistical competency often reflects a minimal portion of formal training [32]. Our results suggest that reliance on traditional models in the face of assumption violations can dramatically skew results toward false positives. This can be attenuated by adopting analyses built for interdependencies. Some of these more complex models were historically impractical for various reasons (e.g., computational intensity, inaccessibility of software), but methods are now more accessible [33]. However, these models can be underpowered at common preclinical sample sizes (Figure 4B). When dealing with categorical data such as those presented here (2+ levels/categories), another model specification that impacts interpretation of results is the coding of categorical variables [34]. Dummy coding sets one level as the “reference,” and results are interpreted in relation to the reference level. This may be relevant when one is interested in changes *from* a given category: optimal decisions in the case of the present data. By contrast, effect coding allows different weights to be assigned to each level. This can allow testing more specific hypotheses (e.g., level 1 vs. level 2 or level 1 vs. average of other levels) and affects the intercept value and its interpretation (effect coding: grand mean across all levels; dummy coding: value of reference level; see supplement S1 for additional details on mixed-effects models).

A key concern for preclinical research is obtaining sufficient power in analyses to detect true effects (i.e., avoid false negatives or Type II error). Animal researchers must either accept that low-N animal studies will inherently have low power or find ways to increase power. These issues have been recently addressed when considering appropriate power and design to include both sexes in preclinical studies [35, 36]. However, there are only a few options to increase power: adjust *p* value threshold, increase effect size, or increase sample size. Increasing the *p* threshold is typically not desirable due to increased false positive rates. Increasing effect sizes may be desirable, but impractical. In pharmacology, a larger effect size might be obtained by increasing drug dose, however, not all drugs have a linear dose-response [37, 38]. A biological manipulation (e.g., lesion/inactivation of a brain region) may lose relevance by making it larger or less specific. This leaves researchers primarily with the option of increasing sample size which has ethical and financial implications, particularly in an era where funding success rates have declined NIH-wide [39].

An alternative to purely increasing power is to increase sensitivity to detect effects. This can be accomplished methodologically, with both statistical analyses and laboratory assay refinement. Power can by increased in a Bayesian analysis by applying priors from historical data to control subjects [19]. In the current study, priors robustly increased power (Figure 5), and thus, our recommendation is to include priors and to control false positives by adopting analyses that account for interdependencies. However, it should be noted that Bayesian methods are not without their drawbacks. In the current study, we limited our analyses (1600 total models) due to extensive computation time. It is more reasonable to perform Bayesian models for the relatively fewer analyses in an experimental study; however, computation time may grow prohibitively long for complex models (e.g., repeated measures, multinomial models) and multiple comparisons. Very strong priors, particularly if they are used for the experimental manipulation, may lend toward false positives when combined with significance testing. This is because as the prior distribution becomes more strongly defined, the observed data has less influence on the ultimate conclusion, particularly when analyzing small sample sizes [40]. Thus, researchers should be cautious in how these tests are applied to their datasets and how priors are obtained. In our data, using the exact prior distribution from real-world data, including narrow error (“strong” priors) started to bias toward effect detection (Figure 5D) and would require a substantial experimental sample size to overcome these effects. Moreover, if the priors obtained are unreliable, this may bias novel analyses. Ultimately, researchers should be aware that these methods will not supplant well-designed and well-powered studies, but instead may be useful to increase power in situations where reliable historical data is available. However, even flat priors improved power over the comparable model (Experiment 3, Figure 4 M4 vs. Figure 5), but potentially at the cost of increased false positive rates (Figure 5A, Figure 6A).

When priors are not feasible (e.g., no historical data available), other approaches may be considered. Statistically, researchers can select a level of analysis at the action or trial level using binomial or multinomial logistic regressions (e.g., Figure 4: M6, M7 in the current data) rather than a session-wide level to gain more observations. Or, if permitted by the experimental design, the actual number of observations can be increased (e.g., trials or sessions). However, this approach may not be feasible for certain designs, such as acute pharmacological challenges or permanent pre/post interventions. We simulated 10 sessions of data for the current study, and the exact effects of session number on power for interdependent data remain unknown, although another published simulation found that 10-25 observations (e.g., trials, sessions) nested within at least 7 clusters (e.g., subjects) were necessary to prevent biased parameter estimates in mixed-effects logistic regression [41]. For a behavioral assessment, researchers may also be able to adjust parameters to gain more sensitivity. This may not work for naturalistic behaviors (e.g., forced swim, elevated plus maze), but trained tasks may have parameters that can be titrated. As an example, a probabilistic reversal task [42] commonly set to 80% vs. 20% for “correct” and “incorrect” choices could be made more difficult by changing to 70% vs. 30% to isolate a more subtle manipulation effect. Moreover, further complications to design occur when considering changes over time. The current simulations were based on stable, steady-state data and thus no within- or between-session effect of learning was modeled. Such learning-oriented data introduce another type of dependency: how a subject’s past behavior predicts their future behavior. These parameters can be accounted for in the models we describe or in more sophisticated time-series, neural network [43], or reinforcement learning models [22], but a high number of observations may be needed for reliable estimation of effects. Additional simulations will be needed to better understand how these factors affect power of such analyses.

The current findings are also of interest to other types of neuroscience data. We used a behavioral example which would immediately apply to any other tasks in which the animal has multiple options about where to go or what to do (e.g., novel object, conditioned place preference, elevated plus maze, water mazes). Monte Carlo simulations for two-option behavioral data already show that accounting for dependency with mixed effects logistic regression is optimal [33, 44], but here we extend this to show that accounting for interdependencies is critical for behavioral data with more than two options. Furthermore, interdependent outcomes also occur in other areas of neuroscience, such as sleep (REM vs. other stages) and relative abundance (e.g., microbiome bacterial composition, cell sequencing clusters). However, paradigms where researchers are interested in interdependent *continuous* outcomes present a more complex problem. For example, time spent swimming, escaping, versus floating on the FST would be more difficult to analyze using logistic regression and likely require binning. However, another option is to use a Bayesian multivariate regression model which analyzes concurrent correlated behaviors.

One final consideration is significance testing methods. Results vary by significance-testing method, particularly for mixed models [45]. Our re-analysis of published data showed that we had not drastically altered the interpretation but did have several parameters incorrectly reported (see Supplement S1). For nested data, there is no agreed-upon method for calculating degrees of freedom and *p*-values. Common methods in R (e.g., Anova() function in the *car* library) often use Wald *z*-tests with Kenward and Roger [46] degrees of freedom estimation for *p*-values. When analyzing large datasets with little correlation among predictions, this approach may be reliable. However, for smaller datasets or correlated predictors, the Wald test can differ from other methods [47]. Other recommendations include likelihood ratio tests, Markov Chain Monte Carlo (MCMC) sampling, and bootstrapping [45, 48]. Here, we used likelihood ratio testing due to computation constraints. Another strategy is to avoid arbitrary cutoffs altogether. This can be accomplished through Bayesian analyses, which generate the likelihood of a range of test statistics (called a credible interval) and allows researcher to present the *strength of evidence for* or *against* an effect.

Our core findings demonstrated that nested choice data was poorly described by traditional linear models. Despite literature demonstrating advantages of GLMs for data bound between 0 and 1 (e.g., percent choice data) [33, 44, 49], linear models on aggregate, interdependent data remain prevalent. A cultural shift is needed in the neurosciences regarding statistics; adoption of modern techniques is a necessity (for a thorough review, see [18]). Neuroscientists are often eager to adopt other advanced techniques (e.g., optogenetics, “-omics” methods, microscopy innovations) and need to approach statistical techniques with similar vigor. Recent changes in NIH policy on transparency in science [50] and journal requirements regarding code and data sharing may help to propel such change, but fundamental shifts in neuroscience education are also needed. Expanded training focused on data literacy and conceptual-level understanding of statistics would enable neuroscientists an appropriate framework from which to learn individual statistical methods and reduce reliance on inappropriate models.

### Conclusions

Poor replication and translation are some of the most consequential problems in preclinical science. The use of inappropriate statistics has likely produced false results, particularly for interdependent data. We showed via simulation that choice data required analyses built to handle interdependencies. Moving forward, statistical methods must be selected based on evidence (e.g., meeting assumptions, accuracy determined via simulation), rather than solely by tradition. Problems with low statistical power can be improved using Bayesian techniques. Dissemination of these practices presents a more complex issue and will require individual and systemic changes.

## Acknowledgements

We would like to thank the researchers in the Vonder Haar laboratory and the Winstanley laboratory who helped to collect the original data which were used to support the simulations in this manuscript.

## Author Contributions

Conceptualization: MF, CV; Formal analysis: MF, PM, MY; Funding acquisition: CV; Writing – first draft: MF; Writing – review and editing: MF, PM, CV, MY

## Data Availability Statement

Code for this project will be shared on the corresponding author’s GitHub repository (https://github.com/VonderHaarLab) at the time of publication and is also included as Supplement S2. Underlying data for TBI simulations (control set) is publicly available at https://odc-tbi.org/, dataset identifier 703. Data for cue manipulation simulations (validation set) was obtained and used with permission from Dr. Catharine Winstanley at the University of British Columbia.

