## Supplement S1 for "Statistical power and false positive rates for interdependent outcomes are strongly influenced by test type: Implications for behavioral neuroscience"

**Additional Resources**

- An interactive, user-friendly tool that provides more information on how the prior distribution affects posterior parameter estimates (Depaoli, Winter, & Visser, 2020): <https://ucmquantpsych.shinyapps.io/sensitivityanalysis/>.
- A tool for calculating sample size and power under Bayesian analyses (Bonapersona, Hoijtink, Sarabdjitsingh, & Joëls, 2021): <https://utrecht-university.shinyapps.io/repair/>. Note that this only allows pairwise calculations at the time of this writing.

**Additional Analyses**

To determine whether previously published data may have been incorrectly reported due to high false positives (see Experiment 1), we re-analyzed our published data using Bayesian methods. Specifically, weak priors from Experiment 3 were used on a mixed-effects regression with Option as a random effect. These were then compared against the published effects (see Table S1). The full code is available as an additional file (S3_Published_Reanalysis.R).

Ultimately, overall effects were not drastically changed. However, the Bayesian analysis appeared more sensitive to detecting effects over time. In the first cohort (Shaver et al., 2019), additional Injury x Time effects were detected. In the second cohort (Shaver et al., 2019), additional Injury x Time effects were also detected, and this rendered the main/initial effects of TBI non-significant across all options. This may be related to the second cohort being tested in acquisition of RGT behavior (as opposed to measuring change from trained behavior). In the third cohort (Ozga-Hess, Whirtley, O'Hearn, Pechacek, & Vonder Haar, 2020), again, additional Injury x Time effects were detected and the main effect of injury on the P2 option was no longer significant. These findings again suggest that there may be better sensitivity of the Bayesian analysis to changes over time. The final cohort (Frankot et al., 2022) had no changes from the Bayesian analysis.

Ultimately, it appears that our prior data had minor errors in reporting of effects, but that ultimately these do not change the conclusions of those studies (TBI still led to deficits that persisted over time). We will seek a correction to clarify the Shaver et al. paper because the second cohort had more substantial changes.

**Additional Considerations**

**Coding of categorical variables.** One specification that impacts results is the coding of categorical variables (Alkharusi, 2012). Categorical regression predictors with more than two levels must be recoded. One common recoding system is dummy coding, where one level is selected as the “reference,” and results are interpreted in relation to the reference level. To generate results in terms of a different level, the variable must be releveled and a new regression run or an estimate of the marginal means difference generated (e.g., *emmeans* package). By contrast, effect coding allows for different weights to be assigned to each level. When effect coding is used, the intercept can be interpreted as the grand mean across all levels, rather than the value of the reference level.

There are other types of recoding, but the key point is that recoding influences interpretation of the intercept. For example, in a choice paradigm, the intercept could represent baseline choice, average choice, or choice of a single option. In our 4-choice intercept-only model, if we used effect coding, the intercept would always equal exactly 25% (i.e., the grand mean across the four options is equal to total choice (100%) divided by four). Thus, an intercept-only model in this context is equivalent to a fixed model. This is particularly important because although fixed predictors default to dummy coding in R, the random effects term in *lme4* defaults to effect coding. Thus, without the addition of *(0+dummy(Option, "2*") to our code, the intercept-only model actually ran as a fixed model. This type of error underscores the importance of proper expertise when using statistical software. When we used dummy coding, the random intercept represented preference for the reference level, optimal choice. Although this approach did indeed vary preference for optimal choice by subject, it resulted in overall preferences that could be greater than or less than 100%. Thus, an intercept-only model is not a good fit for concurrent choice analysis regardless of the recoding technique.

**Analysis of continuous, multivariate outcomes**. It is also notable that we simulated categorical choices. Choice paradigms with continuous outcomes present a more complex problem. For example, time spent swimming, escaping, versus floating on the FST would be more difficult to analyze using logistic regression. Another option is to use a Bayesian multivariate regression model. An analysis of FST data in *brms* could look like the following:

*brm(mvbind(Swim, Climb, Float) ~ TBI + (1|p|Subject), family= “Gaussian”)*

Here, the DV is the three continuous outcomes which are combined using the multivariate binding command *mvbind*. The predictor is *TBI*, and the random effect term varies the intercept by subject. The *|p|* term (symbol is irrelevant – anything can be used) tells the formula to model the random effects as correlated to account for interdependencies (similar to our treatment of *ChoiceOption|Subject* in the main text). The distribution of the variables may also be specified independently with the *family* argument, otherwise a Gaussian distribution is assumed. These subtle model specifications are crucial for accurate analysis, particularly when using flexible programs such as *brms*. More information can be found in the package vignettes in R (*vignette("brms_multivariate")).*

It should be noted that we have not evaluated continuous variables in the current manuscript. The use of these models should be validated prior to use. There may be more challenges applying such analyses to interdependent continuous outcomes compared to categorical.

**Strength and Weakness of Effects in Bayesian Analysis.** A frequentist perspective generally dichotomizes effects as either present or not present (commonly using a *p* value cutoff of 0.05). In contrast, Bayesian analysis presents strength of evidence for or strength against an effect. For example, a 95% credible interval (CI) that has a narrow range and does not overlap with 0 (e.g., 0.5 to 0.6) indicates *strong evidence* for an effect in that range. In contrast, *weak evidence* for the same effect might be a 95% CI which ranges from 0.05 to 1.2. A similar interpretation can be applied to evidence of null effects. For example, a 95% CI with range -0.1 to 0.1 indicates *strong evidence* that there was no effect of this variable. However, if the credible interval ranges from -2 to 2, there is *weak evidence* of no effect. These interpretations can be useful in driving future research. For example, in the case of weak evidence for or against an effect, one might design future experiments with additional controls or manipulations to augment this potential effect and better understand it.

| **Ref** | **Injury** | **Training** | **N**  **(TBI)** |  | **Omnibus** | | **P1** | **P2** | **P3** | **P4** |
| --- | --- | --- | --- | --- | --- | --- | --- | --- | --- | --- |
| Shaver  et al.  2019 | Bilateral  frontal | Trained | 23  (11) | Published  Effect | TBI | *F*(3,3514)  = 7.092  *p* < 0.001 | *t* = 10.14 | *t* = 15.80 | *t* = 6.19 | *t* = 8.96 |
|  |  |  |  |  |  |  | *p* < 0.001 | *p* < 0.001 | *p* < 0.001 | *p* < 0.001 |
|  |  |  |  |  | TBI x  Week |  | *t* = 1.22 | *t* = 3.18 | *t* = 1.79 | *t* = 2.61 |
|  |  |  |  |  |  |  | *p* = 0.224 | *p* = 0.002 | *p* = 0.073 | *p* = 0.009 |
|  |  |  |  | Brms  Effect | TBI |  | β = 1.15 | β = -0.65 | β = 0.85 | β = 0.99 |
|  |  |  |  |  |  |  | CI = (0.65, 1.66) | CI = (-0.93, -0.36) | CI = (0.42, 1.28) | CI = (0.49, 1.48) |
|  |  |  |  |  | TBI x  Week |  | β = -0.24 | β = 0.15 | β = -0.15 | β = -0.24 |
|  |  |  |  |  |  |  | CI = (-0.31, -0.18) | CI = (0.10, 0.20) | CI = (-0.22, -0.08) | CI = (-0.30, -0.17) |
| Shaver  et al.  2019 | Bilateral  frontal | Acquisition | 21  (10) | Published  Effect | TBI | *F*(3,3309)  = 3.66  *p* = 0.012 | *t* = 11.54 | *t* = 21.00 | *t* = 10.82 | *t* = 7.35 |
|  |  |  |  |  |  |  | *p* < 0.001 | *p* < 0.001 | *p* < 0.001 | *p* < 0.001 |
|  |  |  |  |  | TBI x  Week |  | *t* = 0.46 | *t* = 1.37 | *t* = 3.16 | *t* = 0.15 |
|  |  |  |  |  |  |  | *p* = 0.647 | *p* = 0.172 | *p* = 0.002 | *p* = 0.880 |
|  |  |  |  | Brms  Effect | TBI |  | β = 0.57 | β = -0.26 | β = 0.40 | β = 0.16 |
|  |  |  |  |  |  |  | CI = (-0.31, 1.35) | CI = (-0.82, 0.32) | CI = (-0.49, 1.31) | CI = (-0.84, 1.20) |
|  |  |  |  |  | TBI x  Week |  | β = 0.05 | β = -0.07 | β = 0.21 | β = 0.09 |
|  |  |  |  |  |  |  | CI = (-0.03, 0.12) | CI = (-0.12, -0.02) | CI = (0.14, 0.29) | CI = (0.01, 0.17) |
| Ozga-  Hess  et al.  2020 | Unilateral  parietal | Acquisition | 25  (11) | Published  Effect | TBI | *F*(3,3476)  = 16.59  *p* < 0.001 | *t* = 1.68 | *t* = 2.87 | *t* = 1.49 | *t* = 1.50 |
|  |  |  |  |  |  |  | *p* = 0.092 | *p* = 0.004 | *p* = 0.136 | *p* = 0.133 |
|  |  |  |  |  | TBI x  Week |  | *t* = 0.71 | *t* = 4.17 | *t* = 1.08 | *t* = 5.57 |
|  |  |  |  |  |  |  | *p* = 0.480 | *p* < 0.001 | *p* = 0.282 | *p* < 0.001 |
|  |  |  |  | Brms  Effect | TBI |  | β = 0.81 | β = -0.39 | β = 0.15 | β = 0.42 |
|  |  |  |  |  |  |  | CI = (-0.05, 1.69) | CI = (-0.87, 0.09) | CI = (-0.69, 1.02) | CI = (-0.46, 1.34) |
|  |  |  |  |  | TBI x  Week |  | β = 0.13 | β = -0.10 | β = 0.07 | β = 0.24 |
|  |  |  |  |  |  |  | CI = (0.09, 0.17) | CI = (-.13, -0.08) | CI = (0.03, 0.11) | CI = (0.20, 0.28) |
| Frankot  et al.,  2022 | Bilateral  frontal | Trained | 36  (16) | Published  Effect | TBI | No test conducted | *F*(1,31) = 37.32 | *F*(1,31) = 13.83 | *F*(1,31) = 1.55 | *F*(1, 31) = 2.76 |
|  |  |  |  |  |  |  | *p* < 0.001 | *p* < 0.001 | *p* = 0.222 | *p* = 0.107 |
|  |  |  |  |  | TBI x  Week |  | *F*(1,1456) = 0.95 | *F*(1,1456) = 15.89 | *F*(1,1456) = 22.41 | *F*(1,1456) = 10.47 |
|  |  |  |  |  |  |  | *p* = 0.330 | *p* < 0.001 | *p* < 0.001 | *p* = 0.001 |
|  |  |  |  | Brms  Effect | TBI |  | beta = 0.84 | beta = -1.06 | beta = 0.31 | beta = 0.25 |
|  |  |  |  |  |  |  | CI = (0.57, 1.10) | CI = (-1.45, -0.63) | CI = (-0.01, 0.63) | CI = (-0.02, 0.54) |
|  |  |  |  |  | TBI x  Week |  | beta = 0.01 | beta = 0.03 | beta = -0.04 | beta = -0.02 |
|  |  |  |  |  |  |  | CI = (-0.01, 0.02) | CI = (0.02, 0.05) | CI = (-0.05, -0.02) | CI = (-0.03, -0.01) |

**Table S1**: Comparison of published effects against informed Bayesian analysis. Cells with discrepancies are color-coded with their matching cell. A “significant” effect in the Bayesian model was considered a credible interval that did not include 0. In the first cohort (Shaver et al., 2019; Trained), the Bayesian analysis detected significant effects in change over time which were not detected in the original analysis. In the second cohort (Shaver et al., 2019; Acquisition), the Bayesian analysis shifted significance for the P2 and P4 options to change over time rather than the main effect of TBI. It also revealed that reported main effects of TBI on P1 and P3 may have been incorrect. Because this slightly changes the interpretation, an article correction will be requested. In the third cohort (Ozga-Hess et al., 2020), the Bayesian analysis shifted P2 effects to the change over time (large β of 0.10 per week over an 8-week period) and detected significant change over time in the P3 option. In the fourth cohort (Frankot et al., 2022), the Bayesian analysis mapped exactly onto the published analysis. The *emmeans* package was used on the fourth cohort to provide contrasts equivalent to the omnibus/F values reported in the original manuscript.

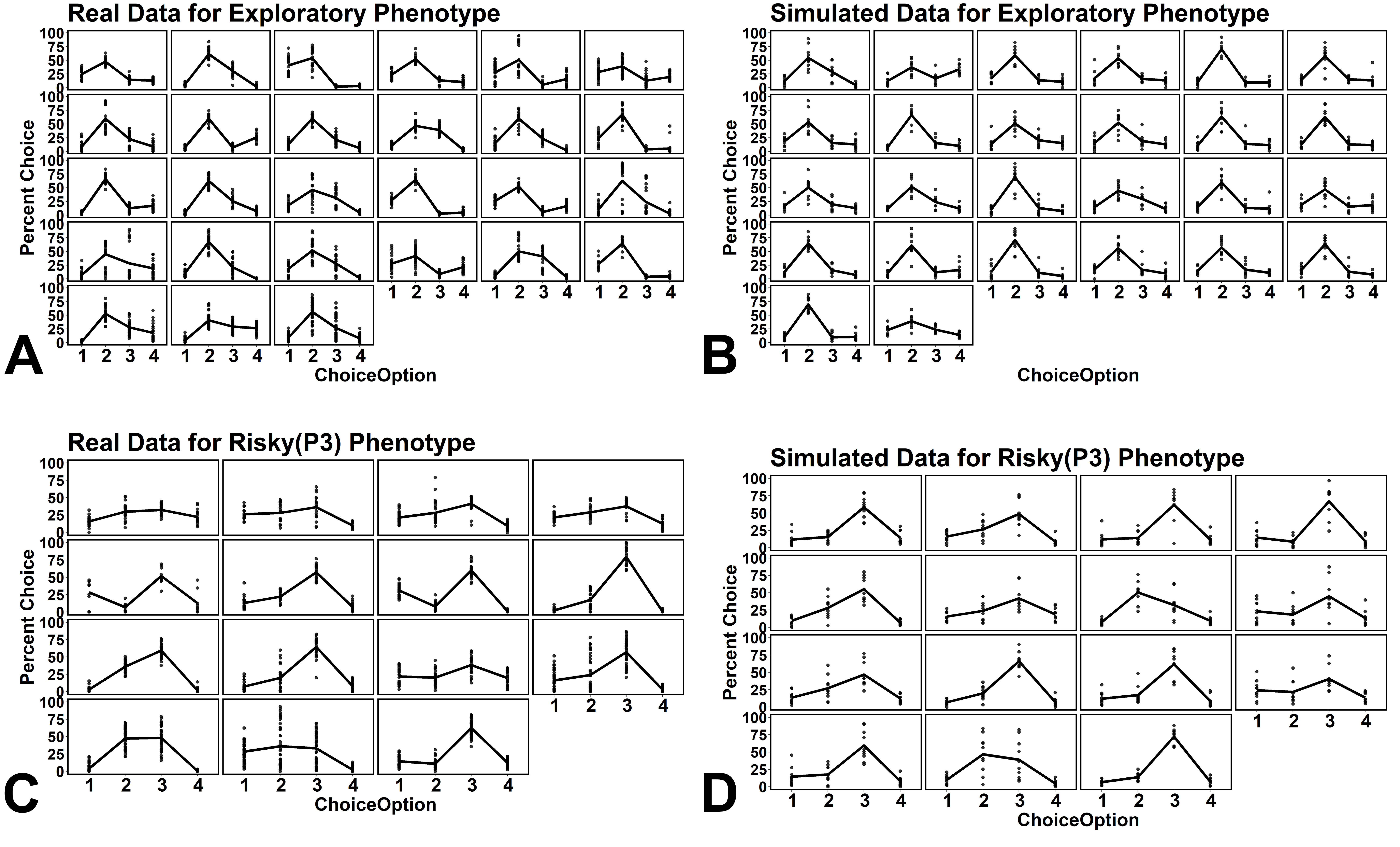

Figure S1. Examples of real collected data (Frankot et al., 2022, Shaver et al., 2020, Ozga-Hess et al., 2020) versus simulated data for the current project. A) The exploratory phenotype. B) Simulated data captures common and rare individuals. C) The risky (P3-preferring) phenotype. D) Simulated data approximates actual individual rats.
