## Supplementary material for "Statistical power and false positive rates for interdependent outcomes are strongly influenced by test type: Implications for behavioral neuroscience": S2: 0b.Simulation.R

```
### Code to simulate RGT data ###

#Conceptual design:
  # In our prior work, we have identified 5 behavioral phenotypes on the RGT
  # We fit an existing data to the softmax function for each phenotype and used those fitted values as population parameters
  # Fitted values are stored in the sim_parameters.csv file
  # The simulation repeatedly samples from those populations, adding variance for each subject and session

#Simulation workflow:
  # Set up: import libraries and file containing simulation parameters
  # Define functions: define functions to vary parameters by subject and session 
  # Simulate data using combination of apply() family and nested loops
  # Process/plot simulated data 

#Note: the parameters are currently set to simulate one dataset, but you can increase the number of repetitions if needed


###set up###

#load libraries 
library(truncnorm) 
library(dplyr)
library(tidyr)
library(readr)
library(progress)


#define softmax function
softmax = function(values, t) {
  exp_values = exp(t*values)
  probs = exp_values/sum(exp_values)
  return(probs) 
}

#import/format reference file for weights/thetas
SM<-read.csv("0a.Simulation_Parameters.csv")
SM<-data.frame(SM)
SM$Phenotype[SM$Phenotype=="p2_high"]<-"p2H" #shorten names for easier reference
SM$Phenotype[SM$Phenotype=="p2_low"]<-"p2L"
row.names(SM)<- paste0(SM$Phenotype, SM$Parameter)
SM['p2HSub_SD', 'P1'] #can reference a single value like this:


####define functions to be used in the simulation###

# First parameter: Theta --> softmax parameter representing how much a subject explores among the RGT choices

#theta by subject function
get_theta_Sub=function(pheno){
  theta_Sub=rtruncnorm(1,a=SM[paste0(pheno,'Sub_min'), 'Theta'], 
                       b=SM[paste0(pheno,'Sub_max'), 'Theta'], 
                       mean=SM[paste0(pheno,'Sub_mean'), 'Theta'], 
                       sd=SM[paste0(pheno,'Sub_SD'), 'Theta'])
  return(theta_Sub)}


#theta by session function
get_theta_Ses=function(Sbj){
  Phenotype_ses<-DF[DF$ID_number==Sbj, "Phenotype"]
  rtruncnorm(1,a=SM[paste0(Phenotype_ses, "Ses_min"), 'Theta'],
             b=SM[paste0(Phenotype_ses,'Ses_max'), 'Theta'], 
             mean=DF$Theta_Sub[DF$ID_number==Sbj],
             sd=SM[paste0(Phenotype_ses,'Ses_SD'), 'Theta'])
}


# Second parameter: Rate --> softmax parameter representing RGT choice preferences


#rate by subject function
get_rate_Sub=function(pheno){
  rate_Sub=c(rtruncnorm(1,a=SM[paste0(pheno,'Sub_min'), 'P1'], b=SM[paste0(pheno,'Sub_max'), 'P1'], 
                        mean=SM[paste0(pheno,'Sub_mean'), 'P1'], 
                        sd=rtruncnorm(1,
                                      a=SM[paste0(pheno,'Sub_SD_min'), 'P1'], 
                                      b=SM[paste0(pheno,'Sub_SD_max'), 'P1'], 
                                      mean=SM[paste0(pheno,'Sub_SD_mean'), 'P1'], 
                                      sd=SM[paste0(pheno,'Sub_SD_SD'), 'P1'])),
             rtruncnorm(1,a=SM[paste0(pheno,'Sub_min'), 'P2'], b=SM[paste0(pheno,'Sub_max'), 'P2'], 
                        mean=SM[paste0(pheno,'Sub_mean'), 'P2'], 
                        sd=rtruncnorm(1,
                                      a=SM[paste0(pheno,'Sub_SD_min'), 'P2'], 
                                      b=SM[paste0(pheno,'Sub_SD_max'), 'P2'], 
                                      mean=SM[paste0(pheno,'Sub_SD_mean'), 'P2'], 
                                      sd=SM[paste0(pheno,'Sub_SD_SD'), 'P2'])),
             rtruncnorm(1,a=SM[paste0(pheno,'Sub_min'), 'P3'], b=SM[paste0(pheno,'Sub_max'), 'P3'], 
                        mean=SM[paste0(pheno,'Sub_mean'), 'P3'], 
                        sd=rtruncnorm(1,
                                      a=SM[paste0(pheno,'Sub_SD_min'), 'P3'], 
                                      b=SM[paste0(pheno,'Sub_SD_max'), 'P3'], 
                                      mean=SM[paste0(pheno,'Sub_SD_mean'), 'P3'], 
                                      sd=SM[paste0(pheno,'Sub_SD_SD'), 'P3'])),
             rtruncnorm(1,a=SM[paste0(pheno,'Sub_min'), 'P4'], b=SM[paste0(pheno,'Sub_max'), 'P4'], 
                        mean=SM[paste0(pheno,'Sub_mean'), 'P4'], 
                        sd=rtruncnorm(1,
                                      a=SM[paste0(pheno,'Sub_SD_min'), 'P4'], 
                                      b=SM[paste0(pheno,'Sub_SD_max'), 'P4'], 
                                      mean=SM[paste0(pheno,'Sub_SD_mean'), 'P4'], 
                                      sd=SM[paste0(pheno,'Sub_SD_SD'), 'P4'])))
  return(rate_Sub)}


#rate by session function
get_rate_Ses=function(Sbj){
  Phenotype_ses<-DF[DF$ID_number==Sbj, "Phenotype"]
  rate_Ses=c(rtruncnorm(1,a=SM[paste0(Phenotype_ses,'Ses_min'), 'P1'], b=SM[paste0(Phenotype_ses,'Ses_max'), 'P1'], 
                        mean=DF$Rate.P1[DF$ID_number==Sbj], 
                        sd=rtruncnorm(1,
                                     a=SM[paste0(Phenotype_ses,'Ses_SD_min'), 'P1'], 
                                     b=SM[paste0(Phenotype_ses,'Ses_SD_max'), 'P1'], 
                                     mean=SM[paste0(Phenotype_ses,'Ses_SD_mean'), 'P1'], 
                                     sd=SM[paste0(Phenotype_ses,'Ses_SD_SD'), 'P1'])),
             rtruncnorm(1,a=SM[paste0(Phenotype_ses,'Ses_min'), 'P2'], b=SM[paste0(Phenotype_ses,'Ses_max'), 'P2'], 
                        mean=DF$Rate.P2[DF$ID_number==Sbj], 
                        sd=rtruncnorm(1,
                                     a= SM[paste0(Phenotype_ses,'Ses_SD_min'), 'P2'], 
                                     b= SM[paste0(Phenotype_ses,'Ses_SD_max'), 'P2'], 
                                     mean= SM[paste0(Phenotype_ses,'Ses_SD_mean'), 'P2'], 
                                     sd= SM[paste0(Phenotype_ses,'Ses_SD_SD'), 'P2'])),
             rtruncnorm(1,a=SM[paste0(Phenotype_ses,'Ses_min'), 'P3'], b=SM[paste0(Phenotype_ses,'Ses_max'), 'P3'], 
                        mean=DF$Rate.P3[DF$ID_number==Sbj],   
                        sd=rtruncnorm(1,
                                     a= SM[paste0(Phenotype_ses,'Ses_SD_min'), 'P3'], 
                                     b= SM[paste0(Phenotype_ses,'Ses_SD_max'), 'P3'], 
                                     mean= SM[paste0(Phenotype_ses,'Ses_SD_mean'), 'P3'], 
                                     sd= SM[paste0(Phenotype_ses,'Ses_SD_SD'), 'P3'])),
             rtruncnorm(1,a=SM[paste0(Phenotype_ses,'Ses_min'), 'P4'], b=SM[paste0(Phenotype_ses,'Ses_max'), 'P4'], 
                        mean=DF$Rate.P4[DF$ID_number==Sbj],  
                        sd=rtruncnorm(1,
                                     a= SM[paste0(Phenotype_ses,'Ses_SD_min'), 'P4'], 
                                     b= SM[paste0(Phenotype_ses,'Ses_SD_max'), 'P4'], 
                                     mean= SM[paste0(Phenotype_ses,'Ses_SD_mean'), 'P4'], 
                                     sd= SM[paste0(Phenotype_ses,'Ses_SD_SD'), 'P4'])))
  return(rate_Ses)}


#third parameter: Trials --> number of trials per session varies across subject/session

#trials by subject function
get_trials_Sub=function(pheno){
  trials_Sub=round(rtruncnorm(1,a=1, b=250, mean=SM[paste0(pheno,'Sub_mean'), 'Trial'], 
                              sd=SM[paste0(pheno,'Sub_SD'), 'Trial']))
  return(trials_Sub)}


#trials by session function
get_trials_Ses=function(Sbj){
  Phenotype_ses<-DF[DF$ID_number==Sbj, "Phenotype"]
  trials_Ses=round(rtruncnorm(1,a=1,b=250,mean=as.integer(DF$Trial_Sub[DF$ID_number==Sbj]),
                              sd=SM[paste0(Phenotype_ses,'Ses_SD'), 'Trial']))
  return(trials_Ses)}


###simulate data using sapply and loop combinations### 

#there are 3 levels of loops 
  # Level 1: assigns subjects to a phenotype and selects population-level softmax parameters
  # Level 2: varies softmax parameters by session 
  # Level 3: assigns choices on trial-by-trial basis

#create dataframes to hold values at all 3 loop levels
DF<-data.frame(Block=double(),Iteration=double(),ID_number=double(), Phenotype=double(), Theta_Sub=double(), 
               Rate.P1=double(), Rate.P2=double(), Rate.P3=double(),Rate.P4=double(), Trial_Sub=double())
DF2<-data.frame(Block=double(),Iteration=double(),Subject=double(), Phenotype=double(), Session=double(),
                Trial_Ses=double(),Theta_Ses=double(), Rate_Ses.P1=double(), Rate_Ses.P2=double(), 
                Rate_Ses.P3=double(),Rate_Ses.P4=double())
DF3<-data.frame(Repetition=double(), Block=double(),Iteration=double(), ID=double(), Phenotype=double(), 
                Session=double(),Trial=double(),ChoiceOption=double())
Counts<-data.frame(ChoiceOption=double(), Session=double(), Subject=double(), 
                       Phenotype=double(), Iteration=double(), ChoiceCount=double())


#Manually change parameters if desired 
num_Block=1 #increase to simulation additional datasets
num_Iter=1 #increase to simulation additional datasets
num_Ses=10
num_Sub=10


#Simulation loop
start<-proc.time() #start timer
pb <- txtProgressBar(min = 0, max = num_Iter*num_Block, style = 3);k<-0 #initiate progress bar
for(Block in 1:num_Block){
  for(Iteration in 1:num_Iter){ 
    k<-k+1;setTxtProgressBar(pb, k);pb #progress bar
    for(ID_number in 1:num_Sub){ #Level 1: assign subjects to phenotype; supply softmax parameters
      Phenotype<-sample(c("p1","p2H", "p2L", "p3", "p4"), 1, replace = TRUE, prob = c(1/58, 33/58, 10/58, 6/58, 8/58)) 
                  #sham probabilities
      #Phenotype<-sample(c("p1","p2H", "p2L", "p3", "p4"), 1, replace = TRUE, prob = c(9/51, 7/51, 17/51, 8/51, 10/51))  
                  #TBI probabilities: effect size estimate = 0.44
                  #un-comment this line to simulate TBI rats instead of Sham rats
      Theta_Sub<-sapply(Phenotype, get_theta_Sub)
      Rate_Sub<-sapply(Phenotype, get_rate_Sub)
      Rate.P1<-Rate_Sub[1];Rate.P2<-Rate_Sub[2];Rate.P3<-Rate_Sub[3];Rate.P4<-Rate_Sub[4]
      Trial_Sub=get_trials_Sub(Phenotype)
      cols<-as.data.frame(cbind(Block, Iteration, ID_number, Phenotype, Theta_Sub, 
                                Rate.P1, Rate.P2, Rate.P3, Rate.P4, Trial_Sub))
      DF=rbind(DF,cols)
    }
    for(Subject in DF$ID_number){ #Level 2: vary softmax parameters by session for each subject
      Theta_Ses<-as.data.frame(matrix(replicate(num_Ses, get_theta_Ses(Subject))));colnames(Theta_Ses)[1] <- "Theta_Ses" 
      Rate_Ses<-as.data.frame(t(replicate(num_Ses, get_rate_Ses(Subject))));colnames(Rate_Ses)=c("P1","P2","P3","P4") 
      Trial_Ses=as.data.frame(matrix(replicate(num_Ses, get_trials_Ses(Subject))));colnames(Trial_Ses)[1] <- "Trial_Ses" 
      Session_df<-as.data.frame(1:num_Ses);colnames(Session_df)[1] <- "Session" 
      Phenotype_df=as.data.frame(rep(DF$Phenotype[DF$ID_number==Subject], num_Ses));colnames(Phenotype_df)[1] <- "Phenotype"
      Iteration_df<-as.data.frame(rep(Iteration,num_Ses));colnames(Iteration_df)[1] <- "Iteration"
      Block_df<-as.data.frame(rep(Block,num_Ses));colnames(Block_df)[1] <- "Block"
      Subject_df<-as.data.frame(rep(Subject,num_Ses));colnames(Subject_df)[1] <- "Subject"
      cols<-as.data.frame(cbind(Block_df,Iteration_df, Subject_df,Phenotype_df, Session_df, Trial_Ses, Theta_Ses, Rate_Ses))
      DF2=rbind(DF2,cols)
    }
    for(Sub in 1:max(as.integer(DF2$Subject))){
      for(Ses in 1:max(as.integer(DF2$Session))){ #Level 3: create trials for each subject/session
        tot_Trial<-as.integer(DF2$Trial_Ses[DF2$Subject==Sub&DF2$Session==Ses])
        ChoiceOption<-replicate(tot_Trial, which.max(rmultinom(1,1,
                                           softmax(c(DF2$P1[DF2$Subject==Sub & DF2$Session==Ses],
                                                     DF2$P2[DF2$Subject==Sub & DF2$Session==Ses], 
                                                     DF2$P3[DF2$Subject==Sub & DF2$Session==Ses], 
                                                     DF2$P4[DF2$Subject==Sub & DF2$Session==Ses]), 
                                                  DF2$Theta_Ses[DF2$Subject==Sub & DF2$Session==Ses]))))
        Iteration_df<-as.data.frame(rep(Iteration,tot_Trial));colnames(Iteration_df)[1] <- "Iteration"
        Block_df<-as.data.frame(rep(Block,tot_Trial));colnames(Block_df)[1] <- "Block"
        Repetition_df<-as.data.frame(Iteration_df$Iteration+((Block_df$Block-1)*num_Iter));colnames(Repetition_df)[1] <- "Repetition"
        Subject_df<-as.data.frame(rep(Sub,tot_Trial));colnames(Subject_df)[1] <- "Subject"
        Session_df<-as.data.frame(rep(Ses, tot_Trial));colnames(Session_df)[1] <- "Session"
        Phenotype_df=as.data.frame(rep(DF2$Phenotype[DF2$Subject==Sub&DF2$Session==Ses], tot_Trial));colnames(Phenotype_df)[1] <- "Phenotype"
        Trial<-as.data.frame(1:tot_Trial);colnames(Trial)[1] <- "Trial"
        cols<-as.data.frame(cbind(Repetition_df,Block_df,Iteration_df,Subject_df, Phenotype_df, Session_df, Trial, ChoiceOption))
        DF3=rbind(DF3,cols)
      }
    }
    #re-intialize dataframes after each iteration
    DF<-data.frame(Block=double(),Iteration=double(),ID_number=double(), Phenotype=double(), Theta_Sub=double(), 
                   Rate.P1=double(), Rate.P2=double(), Rate.P3=double(),Rate.P4=double(), Trial_Sub=double())
    DF2<-data.frame(Block=double(),Iteration=double(),Subject=double(), Phenotype=double(), Session=double(),
                    Trial_Ses=double(),Theta_Ses=double(), Rate_Ses.P1=double(), Rate_Ses.P2=double(), 
                    Rate_Ses.P3=double(),Rate_Ses.P4=double())
  }
  #write raw data to dataframe, process to aggregate data, remove raw data
  count_df<-DF3 %>% count(ChoiceOption, Session, Subject, Phenotype, Repetition) 
  colnames(count_df)[colnames(count_df)=="n"] <- "ChoiceCount"
  Counts<-rbind(Counts, count_df)
  DF3<-data.frame(Repetition=double(), Block=double(),Iteration=double(), ID=double(), Phenotype=double(), 
                  Session=double(),Trial=double(),ChoiceOption=double())
}
proc.time() - start #update timer


###process/plot simulated data###

#assign zeros to implictly missing data
library(tidyverse)
ChoiceSummary<-Counts %>% group_by(Subject, Session, Phenotype, Repetition) %>% 
  complete(ChoiceOption = seq(1,4, by=1), fill = list(ChoiceCount=0)) 


#Calculate percent choice
temp<-aggregate(ChoiceCount~Session+Subject+Repetition, sum, data=ChoiceSummary)
colnames(temp)[colnames(temp)=="ChoiceCount"] <- "TotChoice"
ChoiceSummary<-merge(ChoiceSummary, temp, by=c("Session", "Subject", "Repetition"), all=T)
ChoiceSummary$PctChoice<-(ChoiceSummary$ChoiceCount/ChoiceSummary$TotChoice)*100
ChoiceSummary$ChoiceOption<-as.factor(ChoiceSummary$ChoiceOption)
ChoiceSummary$Session<-as.numeric(ChoiceSummary$Session)
ChoiceSummary$Repetition<-as.numeric(ChoiceSummary$Repetition)


#count number of each phenotype in the simulation
temp<-aggregate(PctChoice~Phenotype+Subject+Repetition, FUN=mean, data=ChoiceSummary)
temp%>%count(Phenotype)


#plotting
library(ggplot2)
my_theme<-theme(
  plot.title = element_text(size=30, face="bold"),
  axis.title.x = element_text(size=25, face="bold"),
  axis.title.y = element_text(size=25, face="bold"),
  axis.text.y = element_text(size=15, face="bold", color="black"),
  axis.text.x = element_text(size=14, hjust = 0, face="bold", color="black"),
  legend.title = element_blank(),
  legend.text = element_text(size = 20, face="bold"),
  legend.key=element_blank(),
  panel.background = element_rect(fill="white", colour="white"),  
  panel.border = element_rect(colour = "black", fill=NA, size=2), 
  strip.text.x = element_text(size = 25, face="bold"),
  strip.text.y = element_text(size = 25, face="bold"),
  strip.background = element_rect(color="white", fill="white"),
  panel.grid.major = element_blank(),
  panel.grid.minor = element_blank(),
) 


#plot choice profiles for individual subjects 
ggplot(data=ChoiceSummary,aes(x=ChoiceOption, y=PctChoice))+   
  geom_point(size=0.8, alpha=0.6)+
  stat_summary(aes(group=Subject), fun=mean, geom="line")+
  facet_wrap(~Subject)+
  ggtitle("Simulation (Sham): Individual Subjects")+
  ylab("Percent Choice")+ 
  my_theme


#plot distributions of each Choice Option
ggplot(ChoiceSummary, aes(x=ChoiceCount)) + 
  geom_histogram(aes(y=..density..), color="darkblue", fill="lightblue", bins=30)+ 
  facet_grid(ChoiceOption~Phenotype)+
  xlab("Choice Count") +
  ylab("Frequency")+
  ggtitle("Simulation (Sham): Choice Distributions")+
  my_theme+
  theme(strip.text.y = element_text(size = 25, face="bold"))


#plot average line for each phenotype
ggplot(data=ChoiceSummary, aes(x=ChoiceOption, y=PctChoice))+   
  geom_point(size=0.8, alpha=0.6)+
  stat_summary(aes(group=Phenotype),size=1, fun=mean, geom="line")+
  facet_wrap(~Phenotype)+
  ggtitle("Simulation (Sham): Choice Profiles")+
  ylab("Percent Choice")+
  my_theme
```
