## Supplementary material for "Statistical power and false positive rates for interdependent outcomes are strongly influenced by test type: Implications for behavioral neuroscience": S2: 1.Exp1-M1-4choice-lm.R

```
### LM for Sham vs. Full TBI in a 4-choice fixed effects linear model
    # Written by Michelle Frankot
    # 6/28/2022
    ### Workflow description: 
        # Step 1: Import/data tidying
        # Step 2: Write function for hypothesis testing
        # Step 3: Run regression in loop
        # Step 4: Calc false positives and false negatives


### Step 1: Import/data tidying ###

#load libraries
library(purrr)
library(readr)


#import Sham data
Sham_df<- read_csv("Sham1000_n20.csv")

#import TBI data (select desired sample size and effect size)
TBI_df<- read_csv("TBI1000_n20_Zero.csv")
TBI_df<- read_csv("TBI1000_n20_0.3.csv")
TBI_df<- read_csv("TBI1000_n20_0.4.csv")
TBI_df<- read_csv("TBI1000_n20_0.5.csv")


#bind data and typcast variables
data<-rbind(TBI_df, Sham_df)
data$Session<-as.numeric(data$Session)
data$Subject<-paste0(data$Subject, data$Injury)
data$Subject<-as.factor(data$Subject)
data$Repetition<-as.numeric(data$Repetition)
data$ChoiceOption<-as.factor(data$ChoiceOption)
data$Phenotype<-as.factor(data$Phenotype)
data$PctChoice<-as.numeric(data$PctChoice)
data$Injury<-as.factor(data$Injury)
data$ChoiceOption<-relevel(data$ChoiceOption, ref="2")


### Step 2. Write function for hypothesis testing ###


  
#function to determine significance via model comparison
get_LMER_aov=function(rep){
  rep_data=subset(data, Repetition==rep)
  m1<-lm(scale(asin(sqrt(PctChoice/100)))~ChoiceOption+Injury,data=rep_data)
  m2<-lm(scale(asin(sqrt(PctChoice/100)))~ChoiceOption*Injury,data=rep_data)
  LMER_aov=as.data.frame(anova(m1, m2, test="Chisq"))
  colnames(LMER_aov) = c('Res.Df', 'RSS', 'Df', 'SS', 'p')
  LMER_aov$Repetition<-rep
  LMER_aov<-LMER_aov[2,]
  return(LMER_aov)}


#plot single dataset
ggplot(data=rep_data, aes(x=ChoiceOption, y=PctChoice, color=Injury))+
  geom_point(alpha=0.7)+
  stat_summary(aes(group=Injury),size=2, fun=mean, geom="line")+
  my_theme
ggplot(data=subset(rep_data, ChoiceOption==2), aes(x=Session, y=PctChoice, color=Injury))+
  geom_point(alpha=0.8)+
  stat_summary(aes(group=Injury),size=2, fun=mean, geom="line")+
  my_theme


### Step 3. Run regressions in loop ###

#initialize data frames
LMER_aov_all=data.frame(Res.Df=double(), RSS=double(), Df=double(), SS=double(), p=double(), Repetition=double())
Errors_all=data.frame(messages=double(),errors=double(), warnings=double(), Repetition=double())
num_Rep=1000
start<-proc.time() #start timer
pb <- txtProgressBar(min = 0, max = num_Rep, style = 3);k<-0 
for(Repetition in 1:num_Rep){
  k<-k+1;setTxtProgressBar(pb, k);pb #progress bar
  #run LMER  using purrr to check for errors; store values in temp_aov for single iteration
  full_LMER <- safely(get_LMER_aov); full_LMER<-quietly(full_LMER); full_model<-full_LMER(Repetition)
  temp_aov=data.frame(Model=double(),Res.Df=double(), RSS=double(), Df=double(), SS=double(), p=double(), Repetition=double()) 
  #if results exist, write to df; if not, write error message
  npar=c(NA);AIC=c(NA);BIC=c(NA);logLik=c(NA);deviance=c(NA);Chisq=c(NA);df=c(NA);p=c(NA);Repetition=Repetition; 
  if(is.null(full_model$result$result)){temp_aov=data.frame(Res.Df, RSS, Df, SS, p, Repetition)
  }else{ temp_aov=rbind(temp_aov, as.data.frame(full_model$result$result))}
  #merge single iteration results with full results
  LMER_aov_all<-merge(temp_aov, LMER_aov_all, 
                      by=c('Res.Df', 'RSS', 'Df', 'SS', 'p', 'Repetition'),all=TRUE)
  #write all errors and messages to dataframe
  Mes_df<-as.data.frame(ifelse(is.null(full_model$messages),NA, full_model$messages))
  Err_df<-as.data.frame(ifelse(is.null(full_model$result$error),NA, full_model$result$error))
  War_df<-as.data.frame(ifelse(is.null(full_model$warnings),NA, full_model$warnings))
  Errors<-cbind(Mes_df, Err_df,War_df,Repetition)
  colnames(Errors) = c('Messages', 'Errors', 'Warnings', 'Repetition')
  Errors_all<-rbind(Errors, Errors_all)
}
proc.time() - start 


### Step 4. calc false positives and false negatives ###


# count results
sum(LMER_aov_all$p <0.05, na.rm=TRUE) #FP
sum(LMER_aov_all$p >0.05, na.rm=TRUE) #FN

#inspect errors
colSums(is.na(Errors_all))


#write data
write.table(LMER_aov_all, "4Choice-fixed_p-vals.csv", sep=",", col.names=NA)
write.table(Errors_all, "4Choice-fixed_n20_Errors.csv", sep=",", col.names=NA)
```
