## Supplementary material for "Statistical power and false positive rates for interdependent outcomes are strongly influenced by test type: Implications for behavioral neuroscience": S2: 7.Exp3-Bayesian.R

```
### Comparison of various Bayesian models for detecting TBI effect in simulated data 
  # Written by Michelle Frankot
  # 7/7/2022
### Workflow description: 
  # Step 1: Import/data tidying
  # Step 2: Write functions for glmer hypothesis-testing
  # Step 3: Run regressions in loop
  # Step 4: Calc false positives and false negatives


### Step 1: Import/data tidying ###

#load libraries
library(purrr)
library(readr)
library(brms) 


#import Sham data
Sham_df<- read_csv("Sham1000_n14.csv")

#import TBI data (one sample size/effect size at a time)
TBI_df<- read_csv("TBI1000_n14_Zero.csv")
TBI_df<- read_csv("TBI1000_n6_0.3.csv")
TBI_df<- read_csv("TBI1000_n14_0.4.csv")
TBI_df<- read_csv("TBI1000_n6_0.5.csv")


#bind data and typcast variables
data<-rbind(TBI_df, Sham_df)
data$Session<-as.numeric(data$Session)
data$Subject<-paste0(data$Subject, data$Injury)
data$Subject<-as.factor(data$Subject)
data$Repetition<-as.numeric(data$Repetition)
data$ChoiceOption<-as.factor(data$ChoiceOption)
data$ChoiceOption<-relevel(data$ChoiceOption, ref="2")
data$Injury<-as.factor(data$Injury)
levels(data$Injury)<-list(TBI  = "TBI_zero", Sham= "Sham")
data$Injury<-relevel(data$Injury, ref="Sham")


### Step 2. Write function for LMER hypothesis-testing ###


#function to perform model comparison between LMERS with and without TBI
calc_model=function(rep){
  rep_data=subset(data, Repetition==rep)
  priors1=c(prior(normal(-0.75, 0.06), class = b,  coef="ChoiceOption1"),
           prior(normal(-0.76, 0.06), class = b,  coef="ChoiceOption3"),
           prior(normal(-0.76, 0.07), class = b,  coef="ChoiceOption4"),
           prior(student_t(3, 1.01, 0.04), class=Intercept)) 
          # prior(normal(0,1),class=b,coef="InjuryTBI")) #need prior on hypothesis to generate BF
  priors2=c(prior(normal(-0.75, 0.2), class = b,  coef="ChoiceOption1"),
            prior(normal(-0.76, 0.2), class = b,  coef="ChoiceOption3"),
            prior(normal(-0.76, 0.2), class = b,  coef="ChoiceOption4"),
            prior(student_t(3, 1.01, 0.15), class=Intercept))
            #prior(normal(0,1),class=b,coef="InjuryTBI"))
  m_prior1 <- brm(asin(sqrt(PctChoice/100)) ~ Injury*ChoiceOption + (ChoiceOption|Subject), 
               data=rep_data,family="Gaussian", chains=4, cores=4, 
               prior=priors1, sample_prior="yes") 
  m_prior2 <- brm(asin(sqrt(PctChoice/100)) ~ Injury*ChoiceOption + (ChoiceOption|Subject), 
                  data=rep_data,family="Gaussian", chains=4, cores=4, 
                  prior=priors2, sample_prior="yes") 
  m_flat <- brm(asin(sqrt(PctChoice/100)) ~ Injury*ChoiceOption + (ChoiceOption|Subject), 
               data=rep_data,family="Gaussian", chains=4, cores=4, 
               sample_prior="yes") 
  mp1<-hypothesis(m_prior1, "InjuryTBI=0"); mp1<- as.data.frame(mp1[["hypothesis"]]); mp1$model="priors1"
  mp2<-hypothesis(m_prior2, "InjuryTBI=0"); mp2<- as.data.frame(mp2[["hypothesis"]]); mp2$model="priors2"
  mf<-hypothesis(m_flat, "InjuryTBI=0"); mf<- as.data.frame(mf[["hypothesis"]]); mf$model="flat"
  results<-rbind(mp1, mp2, mf)
  colnames(results) = c("Hypothesis", "Estimate", "Est.Error","CI.Lower", "CI.Upper", "Evid.Ratio",
                        "Post.Prob", "Star" ,"model")
  results$Repetition<-rep
  return(results)}


### Step 3. Run regressions in loop ###

#initialize data frames
results_all=data.frame(Hypothesis=double(), Estimate=double(), Est.Error=double(), CI.Lower=double(), 
                       CI.Upper=double(), Evid.Ratio=double(), Post.Prob=double(), 
                       Star=double(), model=double(), Repetition=double())
Errors_all=data.frame(messages=double(),errors=double(), warnings=double(), Repetition=double())
num_Rep=100
start<-proc.time() #start timer
pb <- txtProgressBar(min = 0, max = num_Rep, style = 3);k<-0 
for(Repetition in 1:num_Rep){
  k<-k+1;setTxtProgressBar(pb, k);pb #progress bar
  my_analysis <- safely(calc_model); my_analysis<-quietly(my_analysis); analysis<-my_analysis(Repetition)
  temp_results=data.frame(Hypothesis=double(), Estimate=double(), Est.Error=double(), CI.Lower=double(), 
                          CI.Upper=double(), Evid.Ratio=double(), Post.Prob=double(), 
                          Star=double(), model=double(), Repetition=double()) 
  #if results exist, write to df; if not, write error message
  Hypothesis=c(NA);Estimate=c(NA);Est.Error=c(NA);CI.Lower=c(NA);CI.Upper=c(NA)
    Evid.Ratio=c(NA);Post.Prob=c(NA);Star=c(NA);model=c(NA);Repetition=Repetition; 
  if(is.null(analysis$result$result)){
    temp_results=data.frame(Hypothesis, Estimate, Est.Error, CI.Lower, CI.Upper, Evid.Ratio,
                            Post.Prob, Star, model,Repetition)
  }else{
    temp_results=rbind(temp_results, as.data.frame(analysis$result$result))}
  #merge single iteration results with full results
  results_all<-merge(temp_results, results_all, 
                      by=c('Hypothesis', 'Estimate', 'Est.Error', 'CI.Lower', 'CI.Upper', 
                           'Evid.Ratio', 'Post.Prob', 'Star', 'model','Repetition'),all=TRUE)
  #write all errors and messages to dataframe
  Mes_df<-as.data.frame(ifelse(is.null(analysis$messages),NA, analysis$messages))
  Err_df<-as.data.frame(ifelse(is.null(analysis$result$error),NA, analysis$result$error))
  War_df<-as.data.frame(ifelse(is.null(analysis$warnings),NA, analysis$warnings))
  Errors<-cbind(Mes_df, Err_df,War_df,Repetition)
  colnames(Errors) = c('Messages', 'Errors', 'Warnings', 'Repetition')
  Errors_all<-rbind(Errors, Errors_all)
}
proc.time() - start 


### Step 4. calc false positives and false negatives ###


# count results
library(dplyr)
results_all%>%group_by(model)%>%dplyr::count(Star)

#inspect errors
colSums(is.na(Errors_all))
```
