## Supplementary material for "Statistical power and false positive rates for interdependent outcomes are strongly influenced by test type: Implications for behavioral neuroscience": S2: script_pubFigs.R

```
### Plotting Simulation Results ###
      #7/5/2022


#load libraries
library(ggplot2)
library(dplyr)
library(glue)

#set plotting theme
my_theme<-theme(
  plot.title = element_text(size=35, face="bold"),
  axis.title.x = element_text(size=30, face="bold"),
  axis.title.y = element_text(size=30, face="bold"),
  axis.text.y = element_text(size=20, face="bold", color="black"),
  axis.text.x = element_text(size=20, face="bold", color="black"),
  legend.title = element_blank(),
  legend.text = element_text(size = 25, face="bold"),
  legend.key=element_blank(),
  panel.background = element_rect(fill="white", colour="white"),  
  panel.border = element_rect(colour = "black", fill=NA, size=2), 
  strip.text.x = element_text(size = 25, face="bold"),
  strip.text.y = element_text(size = 25, face="bold"),
  strip.background = element_rect(color="white", fill="white"),
  panel.grid.major = element_blank(),
  panel.grid.minor = element_blank(),
) 


#read in data
setwd("~/Desktop/School/Lab/Stats Projects/Sim_Pub/Figures")
data<-read.csv("Simulation_Results.csv")


#typecast variables
data$Sample<-as.numeric(data$Sample)
data$Effect<-as.factor(data$Effect)
data$Type<-as.factor(data$Type)
data$Analysis<-as.factor(data$Analysis)
data$Rate<-as.numeric(data$Raw)
data$Percent<-as.numeric(data$Raw)/10
data<-subset(data, Analysis!="Expected")
data$f<-data$Effect
data$False<-ifelse(data$Effect==0, "Positives", "Negatives")


#break data into Exp 1 vs 2
Exp1<-subset(data, Analysis=="M1:4-choice-lm"|
               Analysis=="M2:4-choice-lmer-intercept"|
               Analysis=="M3:1-choice-lmer-intercept")
Exp1$Analysis<-droplevels(Exp1$Analysis)
Exp2<-subset(data, Analysis=="M4:4-choice-lmer-slope"|
               Analysis=="M5:Score-lmer-intercept"|
               Analysis=="M6:2-choice-glmer-intercept"|
               Analysis=="M7:4-choice-glmer-intercept")
Exp2$Analysis<-droplevels(Exp2$Analysis)


#plot results for Experiment 1
Exp1.colors = c("#18598C", "#33A7D6", "#535A57")
plot.results <-function(type){
ggplot(data=subset(Exp1, False==type), aes(x=Sample, y=Percent, group=Analysis, color=Analysis, shape=Analysis))+
    geom_line(size=2.5)+
    geom_point(aes(fill=Analysis, shape=Analysis),size=10, colour="black", stroke=1)+
    scale_fill_manual(values = Exp1.colors)+
    scale_color_manual(values = Exp1.colors)+
    scale_shape_manual(values=c(23,22,21))+
    facet_wrap(~f, labeller = purrr::partial(label_both, sep = " = "))+
    xlab("Sample Size")+
    ylab(glue("% False {type}"))+
    ggtitle(glue("False {type}"))+ 
    scale_x_continuous(breaks=c(6,10,14,20), labels=c(6,10,14,20))+ 
    coord_cartesian(ylim=c(0,100))+
    expand_limits(x=c(5,21))+
    my_theme}
plot.results("Positives")
ggsave("Exp1_FP.png", width = 30, height = 20, units = "cm")
plot.results("Negatives")
ggsave("Exp1_FN.png", width = 30, height = 20, units = "cm")

#plot power
temp<-subset(Exp1, False=="Negatives")
temp$Power<-100-temp$Percent
ggplot(data=temp, aes(x=Sample, y=Power, group=Analysis, color=Analysis, shape=Analysis))+
  geom_line(size=2.5)+
  geom_point(aes(fill=Analysis, shape=Analysis),size=10, colour="black", stroke=1)+
  scale_fill_manual(values = Exp1.colors)+
  scale_color_manual(values = Exp1.colors)+
  scale_shape_manual(values=c(23,22,21))+
  facet_wrap(~f, labeller = purrr::partial(label_both, sep = " = "))+
  xlab("Sample Size")+
  ggtitle(glue("Statistical Power"))+ 
  scale_x_continuous(breaks=c(6,10,14,20), labels=c(6,10,14,20))+ 
  coord_cartesian(ylim=c(0,100))+
  expand_limits(x=c(5,21))+
  my_theme
ggsave("Exp1_Power.png", width = 30, height = 20, units = "cm")

#d prime calculations
positives<-subset(Exp1, False=="Positives")
negatives<-subset(Exp1, False=="Negatives")
positives$FA<-positives$Percent
positives<- positives[c(1,5,11)]
sdt_data<-merge(positives, negatives, by=c("Sample", "Analysis"))
sdt_data$hits<-100-sdt_data$Percent

library(psycho)
sdt_data$miss<-100-sdt_data$hits
sdt_data$cor_rej<-100-sdt_data$FA
sdt_data<-na.omit(sdt_data)
indices <- psycho::dprime(sdt_data$hits, sdt_data$FA, sdt_data$miss, sdt_data$cor_rej)
sdt_data <- cbind(sdt_data, indices)


#dprime
ggplot(data=sdt_data,aes(x=Sample, y=dprime, group=Analysis, color=Analysis, shape=Analysis))+
  geom_line(size=2.5)+
  geom_point(aes(fill=Analysis, shape=Analysis),size=10, colour="black")+
  scale_fill_manual(values = Exp1.colors)+
  scale_color_manual(values = Exp1.colors)+
  scale_shape_manual(values=c(23,22,21))+
  facet_wrap(~f, labeller = purrr::partial(label_both, sep = " = "))+
  xlab("Sample Size")+
  ylab("d'")+
  ggtitle("Discriminability")+ 
  scale_x_continuous(breaks=c(6,10,14,20), labels=c(6,10,14,20))+ 
  expand_limits(x=c(5,21))+
  my_theme
ggsave("Exp1_dprime.png", width = 30, height = 20, units = "cm")

#c
ggplot(data=sdt_data,aes(x=Sample, y=c, group=Analysis, color=Analysis, shape=Analysis))+
  geom_line(size=2.5)+
  geom_point(aes(fill=Analysis, shape=Analysis),size=10, colour="black")+
  scale_fill_manual(values = Exp1.colors)+
  scale_color_manual(values = Exp1.colors)+
  scale_shape_manual(values=c(23,22,21))+
  facet_wrap(~f, labeller = purrr::partial(label_both, sep = " = "))+
  xlab("Sample Size")+
  ylab("c")+
  ggtitle("Bias")+ 
  scale_x_continuous(breaks=c(6,10,14,20), labels=c(6,10,14,20))+ 
  expand_limits(x=c(5,21))+
  geom_hline(aes(yintercept=0), color="black", linetype="dashed")+
  my_theme
ggsave("Exp1_c.png", width = 30, height = 20, units = "cm")


#plot results for Experiment 2
Exp2.colors = c("#d97575", "#701603", "#5e98eb", "#411052")
plot.results <-function(type){
  ggplot(data=subset(Exp2, False==type), aes(x=Sample, y=Percent, group=Analysis, color=Analysis, shape=Analysis))+
    geom_line(size=2.5, alpha=0.7)+
    geom_point(aes(fill=Analysis, shape=Analysis),size=10, colour="black", alpha=0.9, position=position_jitter(width=0.1, height=0.3))+
    scale_fill_manual(values = Exp2.colors)+
    scale_color_manual(values = Exp2.colors)+
    scale_shape_manual(values=c(23, 22, 21, 24))+
    facet_wrap(~f, labeller = purrr::partial(label_both, sep = " = "))+
    xlab("Sample Size")+
    ylab(glue("% False {type}"))+
    ggtitle(glue("False {type}"))+ 
    scale_x_continuous(breaks=c(6,10,14,20), labels=c(6,10,14,20))+ 
    coord_cartesian(ylim=c(0,10))+
    expand_limits(x=c(5,21))+
    my_theme}
plot.results("Positives")
ggsave("Exp2_FP.png", width = 30, height = 20, units = "cm")

temp<-subset(Exp2, False=="Negatives")
temp$Power<-100-temp$Percent
ggplot(data=temp, aes(x=Sample, y=Power, group=Analysis, color=Analysis, shape=Analysis))+
  geom_line(size=2.5, alpha=0.7)+
  geom_point(aes(fill=Analysis, shape=Analysis),size=10, colour="black", stroke=1, alpha=0.9)+
  scale_fill_manual(values = Exp2.colors)+
  scale_color_manual(values = Exp2.colors)+
  scale_shape_manual(values=c(23, 22, 21, 24))+
  facet_wrap(~f, labeller = purrr::partial(label_both, sep = " = "))+
  xlab("Sample Size")+
  ggtitle(glue("Statistical Power"))+ 
  scale_x_continuous(breaks=c(6,10,14,20), labels=c(6,10,14,20))+ 
  coord_cartesian(ylim=c(0,100))+
  expand_limits(x=c(5,21))+
  my_theme
ggsave("Exp2_Power.png", width = 30, height = 20, units = "cm")

#d prime calculations
positives<-subset(Exp2, False=="Positives")
negatives<-subset(Exp2, False=="Negatives")
positives$FA<-positives$Percent
positives<- positives[c(1,5,11)]
sdt_data<-merge(positives, negatives, by=c("Sample", "Analysis"))
sdt_data$hits<-100-sdt_data$Percent

library(psycho)
sdt_data$miss<-100-sdt_data$hits
sdt_data$cor_rej<-100-sdt_data$FA
sdt_data<-na.omit(sdt_data)
indices <- psycho::dprime(sdt_data$hits, sdt_data$FA, sdt_data$miss, sdt_data$cor_rej)
sdt_data <- cbind(sdt_data, indices)


#dprime
ggplot(data=sdt_data,aes(x=Sample, y=dprime, group=Analysis, color=Analysis, shape=Analysis))+
  geom_line(size=2.5, alpha=0.7)+
  geom_point(aes(fill=Analysis, shape=Analysis),size=10, colour="black", alpha=0.9)+
  scale_fill_manual(values = Exp2.colors)+
  scale_color_manual(values = Exp2.colors)+
  scale_shape_manual(values=c(23, 22, 21, 24))+
  facet_wrap(~f, labeller = purrr::partial(label_both, sep = " = "))+
  xlab("Sample Size")+
  ylab("d'")+
  ggtitle("Discriminability")+ 
  scale_x_continuous(breaks=c(6,10,14,20), labels=c(6,10,14,20))+ 
  expand_limits(x=c(5,21))+
  my_theme
ggsave("Exp2_dprime.png", width = 30, height = 20, units = "cm")

#c
ggplot(data=sdt_data,aes(x=Sample, y=c, group=Analysis, color=Analysis, shape=Analysis))+
  geom_line(size=2.5, alpha=0.7)+
  geom_point(aes(fill=Analysis, shape=Analysis),size=10, colour="black", alpha=0.9)+
  scale_fill_manual(values = Exp2.colors)+
  scale_color_manual(values = Exp2.colors)+
  scale_shape_manual(values=c(23, 22, 21, 24))+
  facet_wrap(~f, labeller = purrr::partial(label_both, sep = " = "))+
  xlab("Sample Size")+
  ylab("c")+
  ggtitle("Bias")+ 
  scale_x_continuous(breaks=c(6,10,14,20), labels=c(6,10,14,20))+ 
  expand_limits(x=c(5,21))+
  geom_hline(aes(yintercept=0), color="black", linetype="dashed")+
  my_theme
ggsave("Exp2_c.png", width = 30, height = 20, units = "cm")


#Experiment 3
Exp3<-read.csv("brms results.csv")
Exp3$Model<-dplyr::recode_factor(Exp3$Model, Flat = "Flat Priors", 
              Weak = "Weakly Informed Priors", Strong = "Strongly Informed Priors")
Exp3.colors<-c("#3f4540", "#407a49", "#92d19c")

#FP
ggplot(data=Exp3, aes(x=as.factor(Sample), y=FP, fill=Model))+
  geom_bar(stat="identity", color="black", position="dodge")+
  ggtitle("False Positives")+
  ylab("% False Positive")+
  xlab("Sample Size")+
  scale_fill_manual(values = Exp3.colors)+
  my_theme
ggsave("Exp3_FP.png", width = 30, height = 20, units = "cm")
  
#FN
ggplot(data=Exp3, aes(x=as.factor(Sample), y=Power, fill=Model))+
  geom_bar(stat="identity", color="black", position="dodge")+
  ggtitle("Statistical Power")+
  ylab("Power")+
  xlab("Sample Size")+
  scale_fill_manual(values = Exp3.colors)+
  my_theme
ggsave("Exp3_Power.png", width = 30, height = 20, units = "cm")


#d prime calculations
library(psycho)
sdt_data<-Exp3
sdt_data$miss<-100-sdt_data$Power
sdt_data$cor_rej<-100-sdt_data$FP
sdt_data<-na.omit(sdt_data)
indices <- psycho::dprime(sdt_data$Power, sdt_data$FP, sdt_data$miss, sdt_data$cor_rej)
sdt_data <- cbind(sdt_data, indices)

ggplot(data=sdt_data, aes(x=as.factor(Sample), y=dprime, fill=Model))+
  geom_bar(stat="identity", color="black", position="dodge")+
  ggtitle("Discriminability")+
  ylab("d'")+
  xlab("Sample Size")+
  scale_fill_manual(values = Exp3.colors)+
  my_theme
ggsave("Exp3_dprime.png", width = 30, height = 20, units = "cm")

ggplot(data=sdt_data, aes(x=as.factor(Sample), y=c, fill=Model))+
  geom_bar(stat="identity", color="black", position="dodge")+
  ggtitle("Bias")+
  ylab("c")+
  xlab("Sample Size")+
  scale_fill_manual(values = Exp3.colors)+
  my_theme
ggsave("Exp3_c.png", width = 30, height = 20, units = "cm")
```
