## Supplementary material for "Statistical power and false positive rates for interdependent outcomes are strongly influenced by test type: Implications for behavioral neuroscience": S2: readme.txt

These files are associated with Frankot et al., 2022, Statistical power 
	and false positive rates for interdependent outcomes are strongly 
	influenced by test type: Implications for behavioral neuroscience.
The current zip archive contains code for generating novel simulated data as described in the manuscript.
	Data which served as the basis for these simulations are publicly available at https://odc-tbi.org/data/703.
A folder with the data generated for the publication is also provided as is syntax for the figures.
R files are provided in the order they are referenced in the publication for the analyses.
