## Supplementary material for "Statistical power and false positive rates for interdependent outcomes are strongly influenced by test type: Implications for behavioral neuroscience": S3

### Supplemental Analyses for Frankot, Young and Vonder Haar, Model Comparison Paper ###
#Re-analysis of existing, published data with methods that account for dependency in data

#Step 1: Download published data from publicly-available dataset & process for analysis
#Step 2: Evaluate data from Shaver et al., 2019, Brain Research (CCI-A study identifier)
#Step 3: Evaluate data from Ozga-Hess et al., 2020, Experimental Neurology (CCI-E study identifier)
#Step 4: Evaluate data from Frankot et al., 2022, Behavioral Neuroscience (HFD-A study identifier)

library(brms)


###Step 1: Download published data from publicly-available dataset & process for analysis

#Data file may be downloaded from: https://odc-tbi.org/data/703
  #This requires creation of an account but is freely available
#Read in data
  data<-read.csv("odc-tbi_703.csv", na.strings="")
#Initial data cleaning
  data<-data[,-c(2:4,6)]
  colnames(data)[colnames(data)=="Choice.P1.4."] <- "ChoiceOption"
  data$Study_ID<-as.factor(data$Study_ID)
  data<-subset(data, Study_ID!="CCI-D")
  data<-subset(data, Session>0)
##Process a Dataset for doing choice analyses
library(dplyr)
library(tidyr)
  #Count function tabulates first option by whatever you list
  #Make sure to include whatever grouping variables are relevant to your dataset
    ChoiceSummary<-data %>% count(ChoiceOption, Subject_ID, Session, Week, Injury, Diet, Pre.Injury, Study_ID) 
    colnames(ChoiceSummary)[colnames(ChoiceSummary)=="n"] <- "ChoiceCount"
  #Remove all zero values (prematures + omissions) from this set
    ChoiceSummary<-subset(ChoiceSummary, ChoiceOption!=0)
      #Zeros (e.g., 0 P4 choice) are not included in count() function, so we need to add them back in
      #Make a set of columns all equal to 0 for each option, then we will bring those into row format
      #Datasets with unique variables (HFD, CCI-D) need to be done separately
  #This will create a zero for all options, and we will add the existing data to it for CCI-A & CCI-E
    temp<-aggregate(ChoiceCount~Subject_ID+Session+Week+Injury+Pre.Injury+Study_ID, mean, data=subset(ChoiceSummary, Study_ID!="HFD-A"))
      #to get a lit of unique sessions by subject. Outcome variable (ChoiceCount) is irrelevant
      #Must retain list of all variables to prevent NAs
    temp<-subset(temp, select=-ChoiceCount)		#delete irrelevant mean variable
    temp$P1<-0			#write in zero'd columns
    temp$P2<-0
    temp$P3<-0
    temp$P4<-0
    temp<-gather(temp, ChoiceOption, ChoiceZero, P1:P4, factor_key=T)
      #Will translate from columns to rows, giving a new variable "ChoiceZero" to be merged with other set
    levels(temp$ChoiceOption)<-c(1,2,3,4)
      #correct the P's that were needed for column names
    ChoiceSummary<-merge(ChoiceSummary,temp, by=c("ChoiceOption", "Subject_ID","Session", "Week", "Injury", "Pre.Injury", "Study_ID"), all=T)
      #Merge the two datasets back
  #This will create a zero for all options, and we will add the existing data to it for HFD-A
    temp<-aggregate(ChoiceCount~Subject_ID+Session+Week+Injury+Pre.Injury+Study_ID+Diet, mean, data=subset(ChoiceSummary, Study_ID=="HFD-A"))
    temp<-subset(temp, select=-ChoiceCount)		#delete irrelevant mean variable
    temp$P1<-0			#write in zero'd columns
    temp$P2<-0
    temp$P3<-0
    temp$P4<-0
    temp<-gather(temp, ChoiceOption, ChoiceZero, P1:P4, factor_key=T)
      #Will translate from columns to rows, giving a new variable "ChoiceZero" to be merged with other set
    levels(temp$ChoiceOption)<-c(1,2,3,4)
      #correct the P's that were needed for column names
    ChoiceSummary<-merge(ChoiceSummary,temp, by=c("ChoiceOption", "Subject_ID","Session", "Week", "Injury", "Pre.Injury", "Study_ID", "Diet"), all=T)
      #Merge the two datasets back
  #Adjust the dataframe we've created
    ChoiceSummary$ChoiceCount[is.na(ChoiceSummary$ChoiceCount)]<-0
      #Replace any NAs that were generated by new data with 0s
    ChoiceSummary<-ChoiceSummary[,-c(10:11)]
  #Calculate total choices & merge back in with dataset
    temp<-aggregate(ChoiceCount~Session+Subject_ID+Week, sum, data=ChoiceSummary)
    colnames(temp)[colnames(temp)=="ChoiceCount"] <- "TotChoice"
    ChoiceSummary<-merge(ChoiceSummary, temp, by=c("Session","Subject_ID","Week"), all=T)
  #Convert each choice to percent
    ChoiceSummary$PctChoice<-ChoiceSummary$ChoiceCount/ChoiceSummary$TotChoice*100
    ChoiceSummary$t.Pct<-asin(sqrt(ChoiceSummary$PctChoice/100))
      #Transformation of percent choice data to better normalize the percents
  #Eliminate any low-observation sessions (e.g., less than 8 total choices)
    ChoiceSummary<-subset(ChoiceSummary, TotChoice>7)
  #Convert character/numeric columns
    summary(ChoiceSummary)
    names<-c(2,4:8)
    ChoiceSummary[,names]<-lapply(ChoiceSummary[,names], as.factor)
    ChoiceSummary$Week<-as.numeric(ChoiceSummary$Week)
    ChoiceSummary$ChoiceOption<-relevel(ChoiceSummary$ChoiceOption, ref="2")
  #priors are selected from the entire dataset for sham animals only. The effect of injury is left as a flat prior.
    priors=c(prior(normal(-0.75, 0.2), class = b,  coef="ChoiceOption1"),
             prior(normal(-0.76, 0.2), class = b,  coef="ChoiceOption3"),
             prior(normal(-0.76, 0.2), class = b,  coef="ChoiceOption4"),
             prior(normal(1.01, 0.15), class=Intercept))

    


###Step 2: Evaluate data from Shaver et al., 2019, Brain Research (CCI-A study identifier)
  ##This paper analyzed rats tested in acquisition (Study_ID = "CCI-A_AQ") and a separate
  ##analysis of rats trained to steady state and assessed after injury (Study_ID = "CCI-A_SS")
  ##These analyses were performed with a mixed-effects model equivalent to lmer(PctChoice~ChoiceOption*Injury*Week+(1|Subject_ID))
      #In the current paper, we show that this analysis is susceptible to false positives.
      #Here we re-analyze with a mixed effects model accounting for each choice option in the random effects
      #and with Bayesian priors to improve power

#Subset data for easier analysis
  Acquisition<-subset(ChoiceSummary, Study_ID=="CCI-A_AQ")
  Trained<-subset(ChoiceSummary, Study_ID=="CCI-A_SS")
    #Baseline calc for Trained
      temp<-aggregate(cbind(PctChoice, t.Pct)~Subject_ID+ChoiceOption, data=subset(Trained, Session>=22 & Session<=26), mean)
      colnames(temp)[3:4]<- paste0(colnames(temp)[3:4],".Base")
      Trained<-merge(Trained, temp, by=c("Subject_ID", "ChoiceOption"))
      Trained<-subset(Trained, Week>0)
  
#Brms re-analysis
  AQ_brm <- brm(scale(t.Pct) ~ Injury*scale(Week)*ChoiceOption+(ChoiceOption|Subject_ID), 
               data=Acquisition, family="Gaussian", chains=4, cores=4, prior=priors) 
    #or read in model if it has already been generated
      AQ_brm <- readRDS("AQ_brm.rds")
    summary(AQ_brm)
    saveRDS(AQ_brm, "AQ_brm.rds")
    
  Trained_brm <- brm(scale(t.Pct) ~ Injury*scale(Week)*ChoiceOption+scale(t.Pct.Base)+(ChoiceOption|Subject_ID), 
                  data=Trained, family="Gaussian", chains=4, cores=4, prior=priors) 
    #or read in model if it has already been generated
    Trained_brm <- readRDS("Trained_brm.rds")
    summary(Trained_brm)
    saveRDS(Trained_brm, "Trained_brm.rds")

    
     
###Step 3: Evaluate data from Ozga-Hess et al., 2020, Experimental Neurology (CCI-E study identifier)
  ##This paper analyzed rats tested in acquisition (Study_ID = "CCI-E") 
  ##These analyses were performed as above and we re-analyze as above

#Subset data for easier analysis
  Unilateral<-subset(ChoiceSummary, Study_ID=="CCI-E")
  Unilateral<-subset(Unilateral, Subject_ID!="CCI-E-8")  #single exclusion due to poor injury in original pub
  Unilateral$Week[Unilateral$Session==518]<-5  #one session mapped to wrong week in second cohort.
  Unilateral$Week<-Unilateral$Week-4 #original analysis just had weeks labeled 1-8 instead of corresponding to post-injury 5-12
  
#BRMS re-analysis
  uni_brm <- brm(scale(t.Pct) ~ Injury*Week*ChoiceOption+(ChoiceOption|Subject_ID), 
                data=Unilateral, family="Gaussian", chains=4, cores=4, prior=priors) 
  #or read in model if it has already been generated
    uni_brm <- readRDS("uni_brm.rds")
  summary(uni_brm)
  saveRDS(uni_brm, "uni_brm.rds")
  
  
  
###Step 4: Evaluate data from Frankot et al., 2022, Behavioral Neuroscience (HFD-A study identifier)
  ##This paper analyzed rats tested in acquisition (Study_ID = "HFD-A") 
  ##These analyses were performed as above and we re-analyze as above
  ##Note that this has a diet manipulation which is analyzed in the original paper and thus is replicated here to provide equivalent models
    
#Subset data for easier analysis
  HFD<-subset(ChoiceSummary, Study_ID=="HFD-A")
  #Baseline calc for Trained
  temp<-aggregate(cbind(PctChoice, t.Pct)~Subject_ID+ChoiceOption, data=subset(HFD, Week==-1), mean)
  colnames(temp)[3:4]<- paste0(colnames(temp)[3:4],".Base")
  HFD<-merge(HFD, temp, by=c("Subject_ID", "ChoiceOption"))
  HFD<-subset(HFD, Week>0)
  
  #BRMS re-analysis
  hfd_brm <- brm(scale(t.Pct) ~ Injury*scale(Week)*ChoiceOption*Diet+t.Pct.Base+(ChoiceOption|Subject_ID), 
                 data=HFD, family="Gaussian", chains=4, cores=4, prior=priors) 
  #or read in model if it has already been generated
    hfd_brm <- readRDS("hfd_brm.rds")
  summary(hfd_brm)
  saveRDS(hfd_brm, "hfd_brm.rds")

  #post hocs using emmeans since it is 2x2 design and we are looking at effect of injury
  #closest comparator to paper since paper reported omnibus/F of TBI
  library(emmeans)
  library(rstanarm)
  emmeans(hfd_brm, pairwise~ Injury|ChoiceOption) 
  pairs(emtrends(hfd_brm, ~Injury*ChoiceOption, var = "Week"))
  
